# A biobank-scale test of marginal epistasis reveals genome-wide signals of polygenic epistasis

**DOI:** 10.1101/2023.09.10.557084

**Authors:** Boyang Fu, Ali Pazokitoroudi, Albert Xue, Aakarsh Anand, Prateek Anand, Noah Zaitlen, Sriram Sankararaman

## Abstract

The contribution of epistasis (interactions among genes or genetic variants) to human complex trait variation remains poorly understood. Methods that aim to explicitly identify pairs of genetic variants, usually single nucleotide polymorphisms (SNPs), associated with a trait suffer from low power due to the large number of hypotheses tested while also having to deal with the computational problem of searching over a potentially large number of candidate pairs. An alternate approach involves testing whether a single SNP modulates variation in a trait against a polygenic background. While overcoming the limitation of low power, such tests of polygenic or marginal epistasis (ME) are infeasible on Biobank-scale data where hundreds of thousands of individuals are genotyped over millions of SNPs.

We present a method to test for ME of a SNP on a trait that is applicable to biobank-scale data. We performed extensive simulations to show that our method provides calibrated tests of ME. We applied our method to test for ME at SNPs that are associated with 53 quantitative traits across ≈ 300 K unrelated white British individuals in the UK Biobank (UKBB). Testing 15, 601 trait-loci associations that were significant in GWAS, we identified 16 trait-loci pairs across 12 traits that demonstrate strong evidence of ME signals (p-value 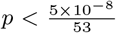). We further partitioned the significant ME signals across the genome to identify 6 trait-loci pairs with evidence of local (within-chromosome) ME while 15 show evidence of distal (cross-chromosome) ME. Across the 16 trait-loci pairs, we document that the proportion of trait variance explained by ME is about 12x as large as that explained by the GWAS effects on average (range: 0.59 to 43.89). Our results show, for the first time, evidence of interaction effects between individual genetic variants and overall polygenic background modulating complex trait variation.

## 1. Introduction

The effect of interactions across genes or genetic variants on a trait (*epistasis*) [1] has been hypothesized to play an important role in human complex trait variation [2, 3]. Understanding the nature and contribution of epistasis is important for elucidating the genetic architecture of complex traits and disease etiology and to improve the accuracy of genetic prediction. Epistasis is one of the factors that could explain missing heritability [4, 5] although some studies suggest a limited contribution of genetic interactions to complex trait variation [6]. Recent studies analyzing the estimates of genetic effects across ancestral populations [7] and the transferability of genetic predictors both within [8] and across ancestries [9, 10] suggest that genetic interactions could explain why genetic effects differ across ancestral populations and the lack of transferability of genetic predictors both within and across ancestries. Epistasis has also been hypothesized to play a role in variable expressivity of complex traits [11]. Nevertheless, our understanding of the role of epistasis in human traits remains limited [12, 5].

Over the past decade, a number of methods to detect epistasis have been developed. The first class of methods explicitly search for pairs of genetic variants (usually single nucleotide polymorphisms or SNPs) that have a non-linear effect on a trait. While allowing for an unbiased search for epistasis (analogous to GWAS enabling an unbiased approach to detect associations), these methods pose serious challenges. Exhaustively searching all pairs of SNPs is computationally difficult (scaling quadratically in the number of SNPs). Further, testing such a large number of hypotheses while controlling the false positive rate requires imposing stringent significance thresholds (scaling quadratically in the number of SNPs if a Bonferroni correction were to be used) which, in turn, reduces power. Efforts to solve this problem have involved the use of statistical techniques [13, 14, 15, 16, 17, 18], algorithmic innovations [19] or hardware infrastructure [20, 21, 22, 23, 24, 25, 26, 27, 28]. Alternate strategies have attempted to reduce the set of SNPs analyzed either restricting to analysis to SNPs identified in GWAS [29, 30, 31] or that are biologically functional [32, 33]. An alternate approach to detect epistasis aims to test for the aggregate epistatic effect across SNPs [34, 35, 36, 37]. Many of these approaches rely on the framework of variance components models that have improved power to detect additive genetic effects in aggregate (in contrast to GWAS that aims to identify individual effects). In this framework, it is of interest to test if the effect of a SNP on a trait is modulated by an individual’s polygenic background. Such tests of *marginal epistasis* [36, 38] can improve power on account of the reduced multiple testing burden (that now scales with the number of SNPs) and due to the aggregation of a number of weak epistatic signals.

Even with the potential improvements in power, it is likely the case that tests of marginal epistasis need to be applied to datasets with large samples to identify robust signals of epistasis [39, 3, 40]. The availability of datasets that contain genetic and phenotypic information across hundreds of thousands of individuals offer an opportunity to detect epistasis with confidence. Estimating marginal epistasis from large data sets such as the UK Biobank consisting of ≈ 500, 000 individuals genotyped at nearly one million SNPs is computationally intractable.

We study the problem of testing whether the effect of a target SNP on a trait is modulated by the genotype of the individual at the remaining SNPs. Given genotypes collected from *N* individuals across *M* SNPs, we consider a model that aims to estimate and test the marginal epistatic effect defined as the combined pairwise interaction effects between a given SNP and all other SNPs while controlling for linear, additive effects. We propose a variance components estimation algorithm to jointly estimate the additive genetic variance component and the marginal epistatic (ME) variance components. The proposed algorithm is efficient both in terms of computation and memory. As a result, our method can be applied to data set with large sample sizes *N* and to estimate the ME of a target SNP to SNPs measured across the genome.

We performed extensive simulations to show that FAME provides calibrated tests of ME and has adequate power to detect true ME signals. We applied FAME to test for ME at trait-associated SNPs for 53 quantitative traits in the UK Biobank (*N* ≈ 300*K* corresponding to unrelated white British individuals and *M* ≈ 500*K* SNPs on the UKBB genotyping array). We explored the robustness of our ME signals to population stratification and to the possibility that the causal variants are missing on the UKBB array. To better characterize the ME signals, we also attempted to partition ME signals to those that fall on the same chromosome and those that fall on the remaining chromosomes as the chromosome containing the target SNP and to estimate the proportion of variance explained by the ME effects.

## 2 Results

### 2.1 Methods overview

We aim to test whether the effect of a target SNP on a phenotype is modulated by the genetic background of the individual by assessing whether the pairwise interactions of the target SNP with each of the remaining SNPs contribute, in aggregate, to variance in the phenotype. In contrast to testing for interactions at a chosen pair of SNPs, this approach of testing for marginal epistasis (ME) can be more powerful when epistatic effects are polygenic, *i*.*e*., we have a substantial number of interactions, each with a weak effect, while also benefiting from the reduced multiple testing burden. To ensure that additive genetic effects are not incorrectly attributed to interactions, we jointly model the additive effects from all genome-wide SNPs (including the target SNP) in addition to the ME effects.

Our method, **FA**st **M**arginal **E**pistasis test (FAME), uses a variance components model in which the phenotypic variance is partitioned into genome-wide additive genetic variance 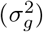, ME variance at a target SNP 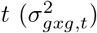, and the residual variance (see Figure 1 for an example and Methods for additional details). The ME variance component 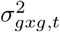 captures the aggregate contribution of all pair-wise interactions between the target SNP *t* and the remaining SNPs in the genome. We would like to test whether the ME variance component is significantly different from zero and, if it is, to be able to estimate its value.

**Figure 1.**
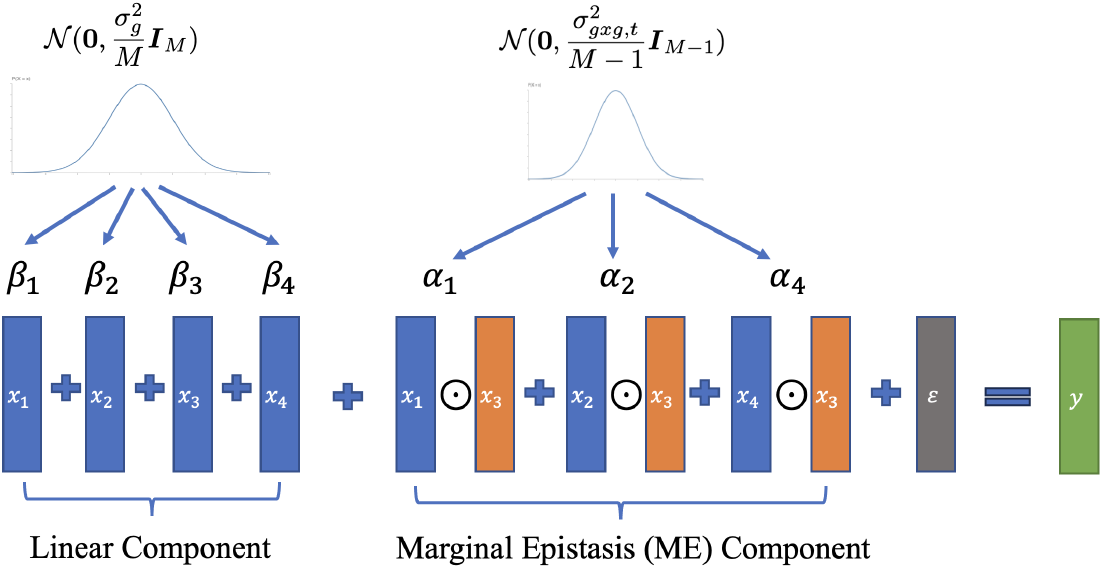
The model underlying FAME. In this example, we have genotypes at four SNPs denoted by **x**_1_, **x**_2_, **x**_3_, **x**_4_. We would like to test for marginal epistasis (ME) between SNP 3 (the target SNP) and the remaining SNPs. We model the relationship of the phenotype ***y*** to the genotypes as arising due to the additive effect of genotypes at each of the four SNPs, the pairwise interaction effects between genotypes at the target SNP (***x***_3_) and the remaining SNPs, and environmental noise ***ϵ***. The additive effect sizes ***β*** are drawn from a distribution with variance parameter proportional to 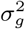 while the ME effect sizes are drawn from a distribution with variance parameter proportional to 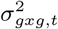 where *t* = 3.

Given a *N × M* genotype matrix ***X*** and a *N* -vector of phenotypes ***y***, we fit the following model [36]:

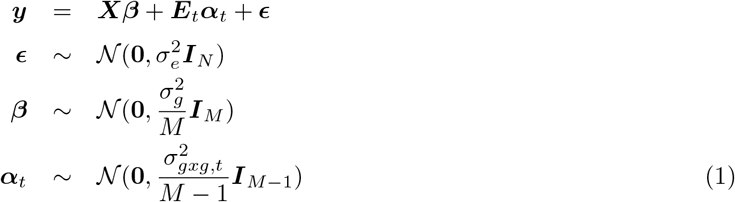

Here 𝒩 (***μ*, Σ**) is a normal distribution with mean ***μ*** and covariance **Σ, *E***_*t*_ denotes a *N ×* (*M* − 1) gene-by-gene interaction matrix defined as ***E***_*t*_ = ***X***_−*t*_ ⊙ ***X***_:*t*_ where ***X***_:*t*_ is the *t*-th column of ***X*** and ***X***_−*t*_ is formed by excluding the column ***X***_:*t*_ from ***X***. In this model, 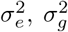, and 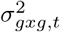 are the residual variance, genetic variance and the ME variance components respectively. ***β*** denotes the additive effects while ***α***_*t*_ denotes the interaction effects between target SNP *t* and each of the other SNPs in the genome. This model assumes that the interaction effects are independent of the main effects so that epistasis is uncoordinated [37].

Fitting this model to Biobank-scale data, containing hundreds of thousands of individuals and millions of SNPs, is computationally impractical. FAME expands on our recent work [41, 42] to be able to test and estimate ME on Biobank-scale data. Specifically, FAME utilizes a randomized Method-of-Moments (MoM) estimator that reduces the size of the input genotype and interaction matrices by multiplying each of these matrices with a pre-specified number (*B*) of random vectors. We show that, even with small values of *B* ≈ 100, this approach results in accurate estimates of the variance components resulting in a highly scalable method.

### 2.2 Calibration of FAME

First, we assessed the false positive rate of FAME by applying it to simulated phenotypes with additive genetic effects but no genetic interactions. We simulated phenotypes based on genotypes from unrelated white British individuals in the UKBB (*M* = 459, 792 SNPs, *N* = 291, 273 individuals). We set the proportion of trait heritability explained by additive genetic effects (additive heritability) 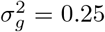 and varied the proportion of variants *p* ∈ {0.01, 0.10} that have non-zero additive effects (causal variants).

The key parameter in applying FAME is the number of random vectors *B* which determines its scalability and stability (see Section 4.2 of Materials and Methods for details). We use *B* = 100 in all our analyses (we explore the impact of this choice in Section 2.5.1). To obtain unbiased estimates, we also do not constrain the estimates of the variance components (allowing for negative estimates). We assessed the calibration of FAME when applied to two sets of target SNPs. The first set consists of target SNPs chosen randomly from across SNPs on the UKBB array. The second set consists of SNPs that were identified to have a significant additive effect based on a GWAS (*p <* 5 *×* 10^−8^) and was chosen to mirror our analyses of traits in UKBB (Section 2.5).

While FAME is calibrated when the target ME SNPs were selected at random (Supplementary Figure S1), the p-values tend to be inflated when the target SNPs were selected based on a GWAS (Section 2.5, Supplementary Figure S2). To address this issue, we excluded SNPs that lie within the LD block around the target SNP when constructing the set of genetic interactions ***E***_*t*_ while retaining these SNPs in the additive component. This approach effectively controlled the false positive rate with no significant ME signal detected across any of the null simulation settings (Figure 2a).

**Figure 2.**
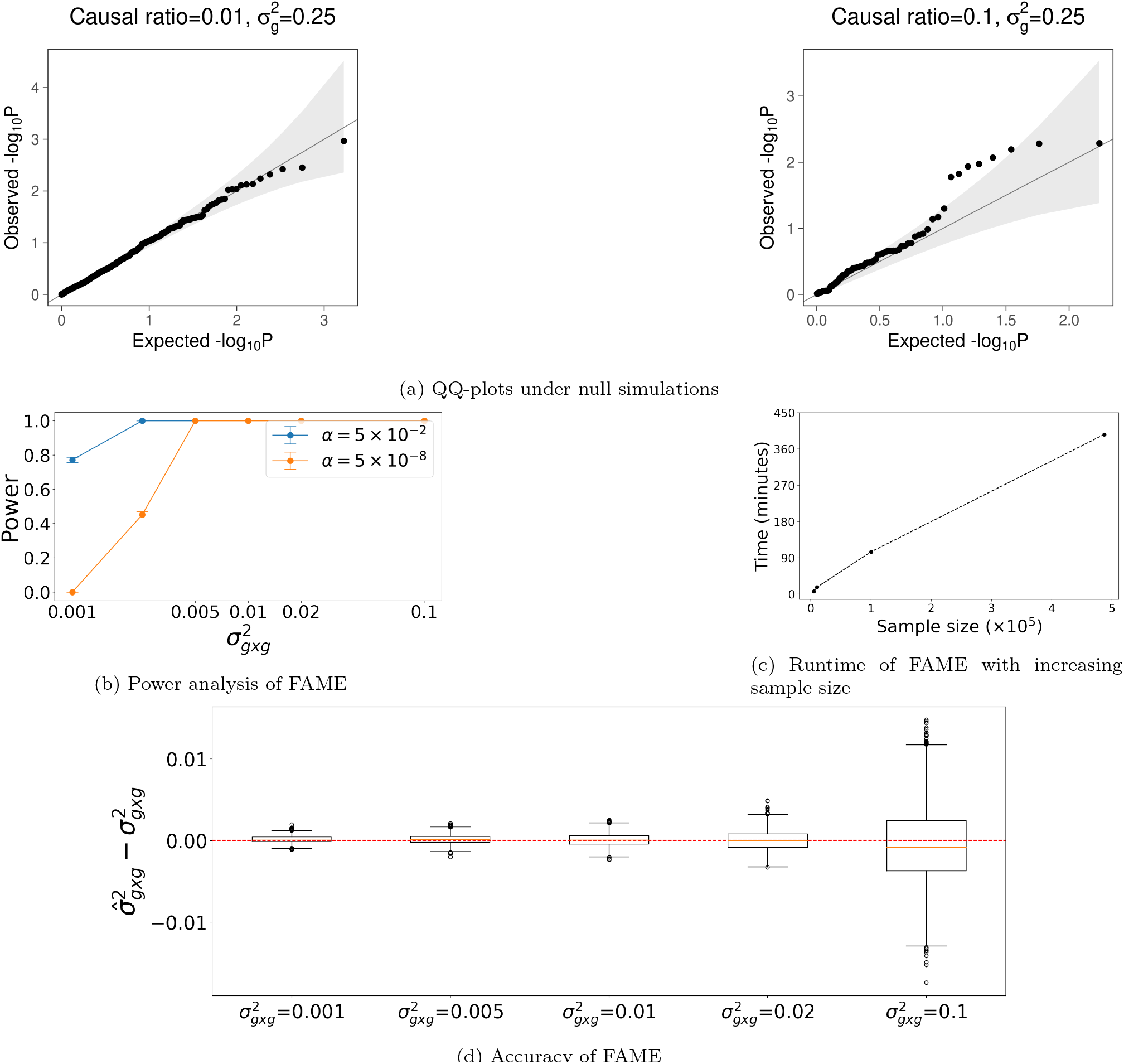
Accuracy and runtime analysis of FAME. (a) QQ-plot of FAME in simulations. We applied FAME to phenotypes simulated from genotypes with linear additive effects but no marginal epistatic (ME) effects. Phenotypes were simulated using genotypes measured on ≈300*K* unrelated white-British individuals in the UK Biobank, with varying ratio of causal SNPs (Causal ratio) and heritability (*h*^2^). We first ran GWAS to identify significant SNPs which were then used as target SNPs in a test of ME. We detected no significant ME signals (*p* ≤ 5 *×* 10^−8^) across all the settings. (b) Power analysis of FAME. We simulated phenotypes by fixing the additive variance component to 0.3 (roughly the median value estimated across real traits). We varied the strength of the ME variance component 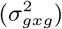. Each setting was simulated 1, 000 times.We plot power for detecting ME at a p-value threshold of 0.05 as well as the genome-wide threshold of 5 *×* 10^−8^ averaged across the replicates. (c). Runtime analysis of FAME. We computed the runtime of FAME applied to common SNPs on the UKBB whole-genome array data and varying sample size. We ran the experiment three times at each setting and reported the average runtime. (d). Accuracy of estimates of the ME variance component 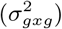 in simulations. We used exactly the same simulation as in (b). We plot the error in the parameter estimates (defined as 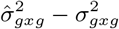) for each parameter setting.

Prior work has shown that tests of epistasis can suffer from inflated false positive rates due to imperfect tagging of causal variants [43, 44, 45]. To examine the robustness of FAME to such imperfect tagging, we repeated the simulations using imputed genotypes (*N* = 291, 273, *M* = 4, 824, 392) while still applying FAME to analyze SNPs on the UKBB array. We observed that FAME remains calibrated indicating its robustness to imperfect tagging (Supplementary Figure S3).

### 2.3 Power analysis

We analyzed the power of FAME by simulating phenotypes with non-zero ME variance components under the model defined in Equation 1. We fixed the variance explained by additive effects, 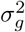 to 0.3, which is roughly the median estimated additive heritability across all the traits we tested. We then varied the proportion of variance explained by ME (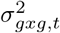 at target SNP *t*) and analyzed the power of FAME to detect ME (see Section 4.3 of Materials and Methods for details). We observed FAME has power ≥ 90% at a stringent p-value threshold (*p <* 5 *×* 10^−8^) even when the variance explained by ME is fairly low 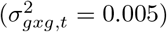 (Figure 2b). Further, we observed that the ME estimates were accurate and exhibited minimal bias (Figure 2d).

### 2.4 Computational efficiency

We attempted to benchmark a previously proposed method for ME testing, MAPIT [36], over datasets of various sizes (see Section 4.4 of Materials and Methods for details). We observe that the computational complexity of MAPIT grows rapidly with increasing sample size even for modest sample sizes and number of SNPs: requiring more than three days to run on 20K samples with 10K SNPs. This is, partly, because MAPIT did not provide the flexibility of modifying the variant testing strategy and, by default, tests the ME effect across all the SNPs provided. Thus, it is not feasible to run MAPIT on a large-scale dataset like UKBB (Supplementary Figure S4). Moreover, MAPIT requires loading the whole genotype matrix at once which we extrapolate would require more than 200 GB for UK Biobank size dataset. On the other hand, FAME can test ME on 500K individuals on a genome-wide dataset containing ≈ 500K SNPs in less than 4 hours (Figure 2c).

### 2.5 Application to UK Biobank phenotypes

We applied FAME to test for ME in 53 quantitative traits measured across *N* = 291, 273 unrelated white British individuals with genotypes measured across common SNPs on the UK Biobank array (see Section 4.5 for details on datasets). Our target SNPs consisted of SNPs that were found to be associated with the trait in a GWAS. Specifically, we ran GWAS on each of the traits, including covariates such as sex, age, and the top 20 genetic PCs. For each trait, we selected SNPs with p-value *p <* 5 *×* 10^−8^ followed by LD pruning (using a window size of 500 SNPs, we computed *r*^2^ between each pair and removed one of them if *r*^2^ *>* 0.1, shifting the window by 1 SNP, and repeating the process). We tested for ME at the resulting set of 15, 601 GWAS significant SNPs in which we also accounted for the linear additive effect of genome-wide SNPs and included age, sex, and the top 20 genetic PCs as fixed effect covariates. Following our calibration experiments, SNPs in the LD block surrounding the target SNP were excluded from the set of genetic interactions. Our tests yielded 21 significant trait-loci pairs across 13 traits (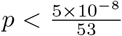 to account for the multiple traits tested). To additionally ensure that the additive genetic effects surrounding the target SNP do not impact estimates of ME, we applied FAME to each of these 21 trait-loci pairs after regressing out all of the SNPs in the LD block of the target SNP to observe 16 trait-loci pairs that retain significant p-values for ME (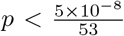 Figure 3a; Table 1).

**Table 1:** Trait-loci pairs with evidence for significant marginal epistasis (ME). *Array* denotes tests of ME run on SNPs genotyped on the UKBB array; *PC40 array* denotes tests performed with the top 40 PCs regressed out instead of top 20 PCs as in the original analysis; *Imputed* represents tests on imputed SNPs. *p* denotes the p-value corresponding to the ME effect while *σ*^2^ denotes the estimate of the ME variance at the tested SNP. p-values that passed the significance threshold 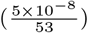 for *PC40 array* and *Imputed* have been highlighted.

| Trait | CHR | SNP | Array |  | *PC40 Array* |  | *Imputed* |  |
| --- | --- | --- | --- | --- | --- | --- | --- | --- |
| | | | $\sigma_{g \times g}^2$<br>$\times 0.01$ | $p$<br>$-\log_{10}$ | $\sigma_{g \times g}^2$<br>$\times 0.01$ | $p$<br>$-\log_{10}$ | $\sigma_{g \times g}^2$<br>$\times 0.01$ | $p$<br>$-\log_{10}$ |
| Alanine aminotransferase | 22 | 44,388,817 | 1.09 | 14.27 | 1.10 | **14.53** | 0.70 | **10.16** |
| Apolipoprotein B | 11 | 116,648,917 | 0.81 | 10.12 | 0.83 | **10.40** | 0.47 | 6.055 |
| C-reactive protein | 1 | 66,257,838 | 0.89 | 9.36 | 0.85 | 8.792 | 0.59 | 6.157 |
| Cholesterol | 11 | 116,648,917 | 0.96 | 13.83 | 0.94 | **13.21** | 0.56 | 8.080 |
| Hemoglobin A1c | 8 | 41,542,093 | 0.65 | 14.72 | 0.66 | **14.97** | 0.58 | **15.05** |
| Lipoprotein-A | 6 | 160,578,069 | 1.42 | 19.24 | 1.41 | **19.22** | 1.50 | **41.33** |
|  |  | 160,560,845 | 1.77 | 17.18 | 1.79 | **17.43** | 1.09 | **13.52** |
|  | 19 | 45,414,399 | 1.15 | 14.75 | 1.14 | **14.61** | 0.77 | **10.04** |
| Mean platelet volume | 20 | 57,597,970 | 0.89 | 18.25 | 0.88 | **17.82** | 0.73 | **16.03** |
| Monocyte count | 13 | 28,623,048 | 0.79 | 23.65 | 0.79 | **23.92** | 0.66 | **21.49** |
| SHBG | 17 | 7,145,117 | 1.06 | 24.41 | 1.09 | **25.12** | 0.91 | **21.80** |
|  |  | 7,254,315 | 0.98 | 15.55 | 0.98 | **15.56** | 0.74 | **12.82** |
| Testosterone | 7 | 99,032,593 | 0.82 | 11.29 | 0.81 | **11.13** | 0.63 | **9.32** |
|  | 12 | 2,977,954 | 0.93 | 25.41 | 0.91 | **24.88** | 0.84 | **23.64** |
| Triglycerides | 11 | 116,648,917 | 1.77 | 11.62 | 1.75 | **11.74** | 1.08 | **16.10** |
| Urate | 1 | 145,630,111 | 0.44 | 14.44 | 0.44 | **14.58** | 0.40 | **14.37** |

**Figure 3.**
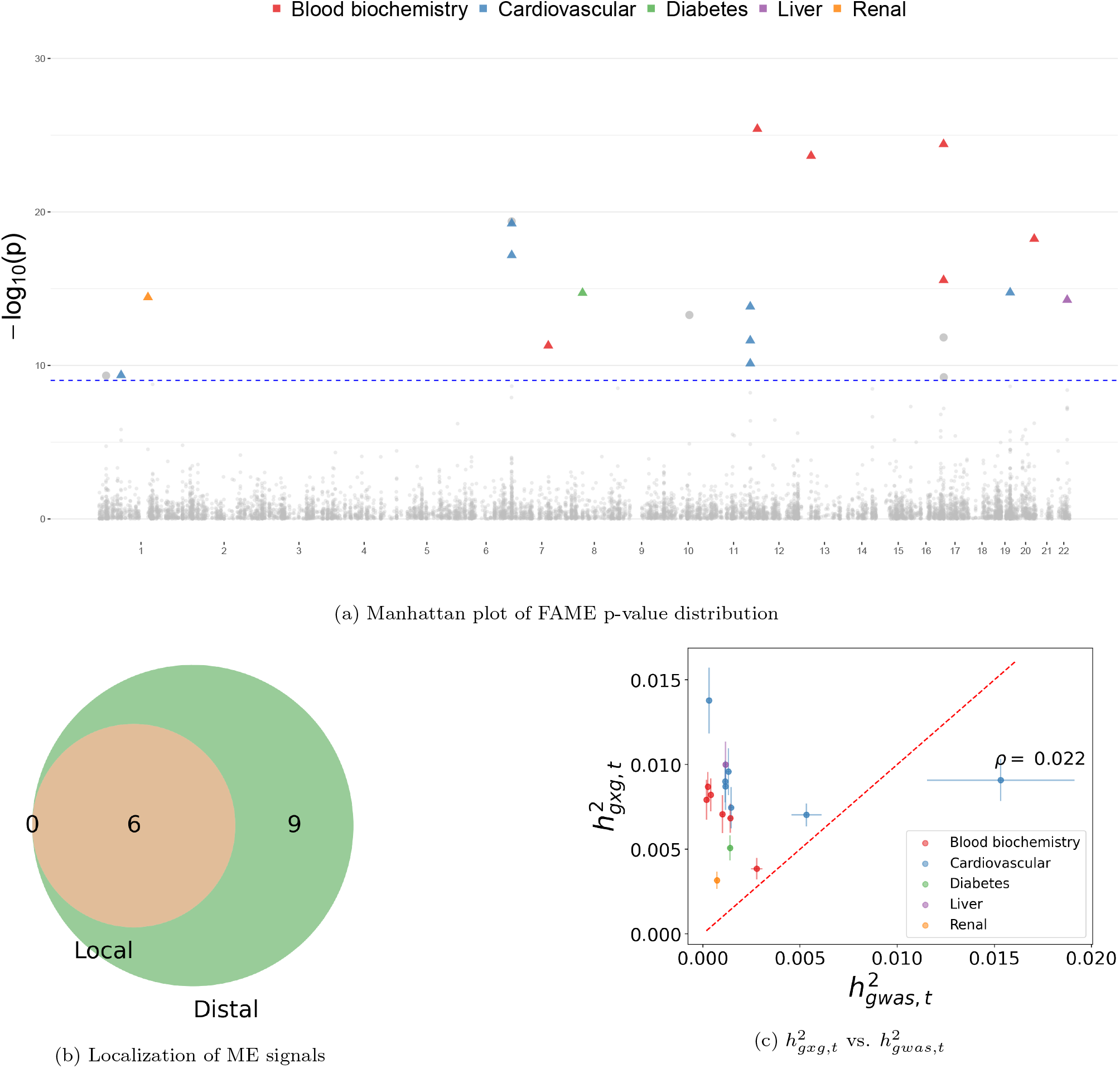
ME signals in the UKBB. (a) Manhattan plot of the ME loci across 53 complex traits in UKBB. Colored shapes denote trait-loci pairs that are significant at 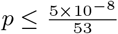 ; shapes with colored triangles were the loci that are statistically significant in our initial analysis and after we regressed out all SNPs within the LD block as fixed effects. (b) Localization of ME signals. For each of 16 trait-loci pairs, we tested whether the ME signals remained significant when testing against all SNPs on the same chromosome as the target SNP (after removing SNPs in the same LD block as the target SNP), which we term *local*, and against all SNPs on chromosomes different from the chromosome containing the target SNP, which we term *distal*. We then compared the overlap between the *local* and *distal* significant signals 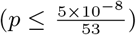. (c) We compared the fraction of phenotypic variance explained by marginal epistatic effects 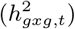 to the fraction of phenotypic variance explained by GWAS (denoted as the 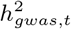) for trait-loci pairs that show significant ME. Vertical (horizontal) bars denote the standard error of 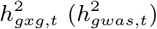.

#### 2.5.1 Stability of significant ME signals

We first explored the impact of the randomization underlying FAME on our results. We selected two traits: body mass index (BMI), for which we did not detect a significant ME locus, and serum urate levels (Urate), for which we detected a significant ME locus. We computed the Pearson correlation of the negative log p-value between results of FAME run with different seeds (*ρ*). We experimented with the number of random vectors (*B*) and observed that using *B* = 100 random vectors yields consistent results (*ρ* = 0.99 for Urate; *ρ* = 0.98 for BMI; Supplementary Figure S6). Second, we reran FAME for the 16 significant trait-loci pairs using five different random number seeds and found that the results are concordant across seeds (15 of the trait-loci pairs show 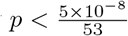 across all seeds while all of the trait-loci pairs show *p <* 5 *×* 10^−8^ across all seeds; Supplementary Table S1). These results indicate that FAME yields stable estimates of ME.

#### 2.5.2 Robustness of significant ME signals

Population stratification in GWAS is commonly accounted for by including principal components (PCs) computed from genotype data as covariates in the analysis [46, 47]. To explore the effect of population stratification, we reran our analyses on trait-loci pairs previously discovered as significant with the number of PCs included as covariates increased to 40 (from 20). We observe a high correlation in the p-values when using 40 vs 20 PCs (Table 1, Supplementary Figure S7a; Pearson correlation *ρ* = 0.997). Importantly, 15 of the 16 significant trait-loci pairs remain significant after including the top 40 PCs, indicating that our findings are robust to population stratification (with the remaining trait-locus pair continuing to exhibit a low p-value).

A second concern with our analyses arises from the fact that the UK Biobank array might miss true causal variants which could lead to the inference of spurious epistatic effects [43, 44, 45]. Our simulations in Section 2.2 show that FAME remains calibrated in this setting. To further explore the robustness of our results, we analyzed our significant ME signals on 4, 824, 392 imputed SNPs (MAF *>* 1%). We observed 13 out of the 16 significant trait-loci pairs detected on the array dataset were significant 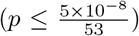 on the imputed dataset (the remaining three loci had p-values *p* ≤ 10^−6^ on the imputed dataset; Table 1; Pearson correlation of the p-values *ρ* = 0.673; Supplementary Figure S7b).

Third, we observe that SNPs with significant ME effects were associated with lower MAF compared to GWAS significant SNPs (Supplementary Table S2). To evaluate the calibration of FAME at low MAF SNPs, we repeated our null simulations described in Section 2.2 restricting to low MAF candidate causal variants (MAF ∈ [0.01, 0.05]). We confirm that FAME remained calibrated across all tested settings (Supplementary Figure S5).

It is well-known that scale of phenotype measurement can affect approaches to test and interpret epistasis. If epistatic effects arose due to choice of scale, we would expect a genome-wide impact for the associated SNPs. However, we do not observe widespread inflation in tests of ME suggesting that the choice of scale is unlikely to impact our results. To further explore the impact of scale, we selected one of the phenotypes (height) and randomly chose 100 GWAS significant SNPs as target SNPs. We then ran FAME by changing the scale of the trait considering two possible transformations: *y*^*pow*3^ := *y*^3^ and *y*^*exp*^ := *exp*(*y*). We then compared the p-value of the ME test to those obtained by analyzing height on the original scale *y*^*orig*^ := *y*. We noticed that by changing the scale of the target trait, the p-values from FAME were significantly inflated (Supplementary Figure S9), thus suggesting that the ME signals discovered by FAME are unlikely to arise due to the scale on which traits are measured.

#### 2.5.3 Localizing signals of ME

Having demonstrated evidence for genome-wide ME, we sought to understand where these interactions localize. As a first step towards answering this question, we extended FAME to test for ME of a target SNP with only a subset of SNPs while accounting for the additive effects of genome-wide SNPs. We separately tested for ME of the target SNP with other SNPs that fall on the same chromosome (local ME; denoted as *gxg*_*local*_) and the ME of the target SNP with SNPs located on chromosomes distinct from the chromosome containing the target SNP (distal ME; denoted as *gxg*_*dist*_). We first confirmed that tests of 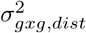 and 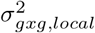 are well-calibrated in simulations (Supplementary Figure S8; see Section 4.3 of Materials and Methods for details). Applying the localization test to each of 16 previously identified ME loci, we found 6 and 15 loci with significant local and distal ME effects respectively (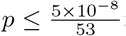 ; Figure 3b; Supplementary Table S3).

#### 2.5.4 Magnitude of ME effects

We estimated the proportion of trait variance explained by ME (ME heritability) at a target SNP 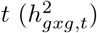 from the variance components estimated by FAME (Supplementary Information Section S1). Across the 16 trait-loci pairs with significant ME signal, estimates of 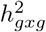 tend to be modest: 10^−3^ − 10^−2^. We compared these estimates to the heritability of the SNP based on its GWAS effect size (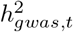 estimated as the square of the GWAS effect size for a standardized genotype). We find that the 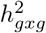 estimates are substantially larger than the corresponding 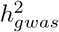 estimates: about 12*x* larger on average with a range of 0.59 to 43.89 (Figure 3c; Table 2). Estimates of 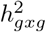 are not strongly correlated with the 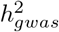 estimates (*ρ* = 0.022).

**Table 2:** Analysis of the heritability at loci with significant ME effects. For each SNP *t* with significant ME, we report estimates of the ME heritability 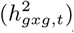, the heritability of the SNP based on its GWAS effect 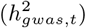, the standard error (SE), and the ratio between 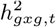 and 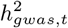. The ME effects were estimated after regressing out the additive effect within the LD region of the target SNPs.

| Trait | CHR | SNP | $h^2_{g_{xg},t}$<br>$\times 0.001$ | $SE(h^2_{g_{xg},t})$<br>$\times 0.001$ | $h^2_{g_{was},t}$<br>$\times 0.001$ | $SE(h^2_{g_{was},t})$<br>$\times 0.001$ | Ratio |
| --- | --- | --- | --- | --- | --- | --- | --- |
| Alanine aminotransferase | 22 | 44,388,817 | 10.003 | 1.352 | 1.168 | 0.080 | 8.562 |
| Apolipoprotein B | 11 | 116,648,917 | 7.457 | 1.218 | 1.451 | 0.111 | 5.140 |
| C-reactive protein | 1 | 66,257,838 | 8.703 | 1.389 | 1.157 | 0.079 | 7.520 |
| Cholesterol | 11 | 116,648,917 | 9.001 | 1.231 | 1.150 | 0.078 | 7.828 |
| Hemoglobin A1c | 8 | 41,542,093 | 5.078 | 0.737 | 1.397 | 0.104 | 3.636 |
| Lipoprotein-A | 6 | 160,560,845 | 13.781 | 1.949 | 0.314 | 0.011 | 43.888 |
|  | 19 | 45,414,399 | 9.587 | 1.389 | 1.315 | 0.095 | 7.291 |
|  | 6 | 160,578,069 | 7.029 | 0.668 | 5.330 | 0.778 | 1.319 |
| Mean platelet volume | 20 | 57,597,970 | 6.830 | 0.864 | 1.421 | 0.107 | 4.805 |
| Monocyte count | 13 | 28,623,048 | 3.851 | 0.628 | 2.771 | 0.292 | 1.390 |
| SHBG | 17 | 7,145,117 | 8.211 | 0.965 | 0.410 | 0.017 | 20.004 |
|  |  | 7,254,315 | 7.064 | 1.126 | 1.006 | 0.064 | 7.021 |
| Testosterone | 7 | 99,032,593 | 7.915 | 1.176 | 0.184 | 0.005 | 42.981 |
|  | 12 | 2,977,954 | 8.687 | 0.865 | 0.260 | 0.008 | 33.473 |
| Triglycerides | 11 | 116,648,917 | 9.083 | 1.242 | 15.326 | 3.795 | 0.593 |
| Urate | 1 | 145,630,111 | 3.166 | 0.495 | 0.720 | 0.039 | 4.395 |

#### 2.5.5 Interpreting loci with significant ME effects

We observe the largest ratio of 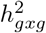 to 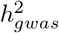 at SNP rs628031 (chr6:160,560,845) that shows significant ME for serum lipoprotein A levels (lipoA). This variant is a non-synonymous polymorphism that changes methionine to valine in the protein product of the organic cation transporter gene OCT1 (also known as SLC22A1). OCT1 mediates the uptake and efflux of cationic metabolites in the liver that includes as its substrates a variety of drugs including metformin that is widely used to treat type 2 diabetes [48]. Genetic variation in OCT1 has been shown to modulate the response to metformin and to other drugs [48].

SNP rs964184 (chr11:116,648,917) shows significant ME for multiple traits: Apolipoprotein B, cholesterol, and triglycerides with substantial ME effects (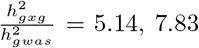, and 0.59 respectively). This variant lies in the 3’ UTR region of the ZPR1 gene (also referred to as ZNF259) that encodes a zinc finger protein that is known to play a regulatory role in cell proliferation and signal transduction [49]. The promoter region of ZPR1 is known to be bound by transcription factors that play a role in insulin sensitivity, cholesterol metabolism, and obesity. rs964184, as well as other variants in ZPR1, have been found to be associated with serum LDL-C [50], HDL-C [51], triglyceride levels [50, 52, 53] and risk for coronary artery disease (CAD) [54] in diverse populations. A regulatory role for rs964184 has been suggested based on its location in a DNaseI hypersensitive region and its overlap with an enhancer that is active in tissues relevant for lipid biology [52]. Further, rs964184 has been association with DNA methylation of a CpG site in the promoter region of the APOA5 gene [55], potentially explaining the association between DNA methylation level at this site and triglyceride levels [56]. Integrative analyses of genotype and gene expression data have shown rs964184 to play a regulatory role: being a cis-eQTL for genes PCSK7, SIDT2, TAGLN, and BUD13 while also a trans-eQTL for TMEM165, YPEL5, PPM1B, and OBFC2A [57]. Further, mediation analyses revealed that a substantial proportion of the effect of rs964184 on HDL-C and triglycerides is mediated through its trans association with PPM1B and YPEL5 [57].

## 3 Discussion

We have presented a new method, FAME, that can detect marginal epistasis (ME) in Biobank-scale data. FAME yields calibrated results in simulations. Applying FAME to 53 quantitative phenotypes in the UK Biobank, we found 16 trait-loci pairs with significant signals of ME, a vast majority of which remain significant after testing with additional PCs to correct for population stratification, and on imputed genotypes to reduce the impact of missing causal SNPs. To the best of our knowledge, this work is the first to show evidence of interaction effects between individual genetic variants and overall polygenic background modulating complex trait variation. While the number of loci showing ME effects is modest (in part due to the stringent p-value threshold that we impose and the GWAS selection strategy that we used to identify target SNPs), we observe that the proportion of variance explained by ME is comparable to, and sometimes substantially larger than, the proportion of variance explained by GWAS. These results show that the polygenic background can substantially modulate the effect of a genetic variant on trait and has implications for efforts to interpret genetic variant effects, to improve phenotype prediction, and to understand how genetic effects vary across populations [7].

We further partitioned the ME signal within and across chromosomes to detect both within and cross-chromosomal signals and found 6 within chromosomal signals, which is a strict subset of the 15 cross-chromosomal signals. This observation suggests that the epistatic signal that we detect is likely to be polygenic so that the approach of testing for the aggregate effects as we do here is likely to be more powerful than an approach that aims to identify specific pairs of SNPs. While our current application of FAME has focused on genome-wide signals of ME where we test a single target SNP against a background set consisting of SNPs across the genome (excluding those in the LD block as the target), the model underlying FAME is flexible and can be applied to test for epistasis in other settings. For example, FAME can be extended to test for interactions of a target SNP or other covariates (such as polygenic scores) with a background set of SNPs where the set is defined based on functional annotation such as genes or pathways. The ideas underlying FAME allow such tests to be applied to biobank-scale data. Such an approach can improve on our understanding by attempting to localize the ME signal. Additionally, the model underlying FAME assumes that epistasis is uncoordinated, *i*.*e*., the interaction effects are independent of main effects. It would be of interest to extend our method to settings where epistasis is coordinated [37].

Our work has several limitations. First, it is plausible that the impact of population structure on epistatic effects might not be well-modeled by the approaches employed here (such as the inclusion of principal components based on common genetic variants). Second, prior studies have shown that tests of epistasis can have inflated false positive rates due to imperfect tagging of causal variants that have large additive effects [43]. Our simulations show that FAME is robust to imperfect tagging of causal variants. Further, the replication of signals discovered using array SNPs on imputed SNPs, that are unlikely to miss causal variants that are common in the population, makes the issue of missing causal variants less likely. Nevertheless, it is plausible that the distributions of causal variants and the LD patterns between causal and genotyped variants could be complex which could impact the calibration of our method. Third, the scale on which phenotypes are measured can affect our results (as is true of other approaches to detect epistasis). Our simulations applying FAME to rescaled versions of phenotypes and the observation that we find SNPs with significant and non-significant ME indicate that our results are not simply driven by scale. Fourth, our estimates of ME effects are likely to be biased upwards due to winner’s curse [58]. Fifth, despite its scalability, FAME is still not efficient enough to perform genome-wide scans of ME which, in turn, led us to focus on testing for ME at GWAS loci. Extending the scope and efficiency of FAME present important directions for future work.

## 4 Materials and Methods

### 4.1 Marginal epistasis model

Given a *N × M* genotype matrix ***X***, a *N* -vector of phenotypes ***y*** and a target SNP *t* ∈ {1, …, *M* }, we aim to jointly test the additive effect of the *M* SNPs and the ME of the target SNP based on the following model that was originally introduced in [36]:

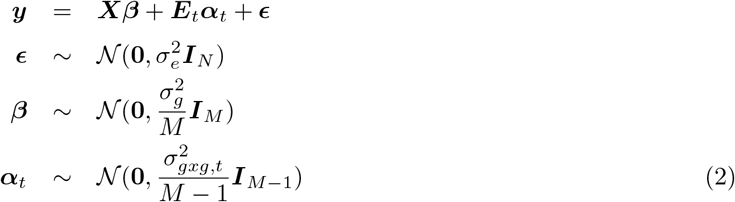

Here 𝒩 (***μ*, Σ**) is a normal distribution with mean ***μ*** and covariance **Σ, *E***_*t*_ denotes a *N ×* (*M* − 1) gene-by-gene interaction matrix defined as ***E***_*t*_ = ***X***_−*t*_ ⊙ ***X***_:*t*_ where ***X***_:*t*_ is the *t*-th column of ***X*** and ***X***_−*t*_ is formed by excluding the column ***X***_:*t*_ from ***X***.

In this model, 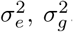, and 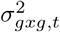 are the residual variance, genetic variance and the ME variance components respectively. ***β*** denotes *M* -vector of SNPs effect sizes and ***α***_*t*_ denotes *M* − 1-vector of interaction effects between target SNP *t* and each of the other SNPs in the genome.

We assume without loss of generality that ***y*** is centered and the columns of ***X*** are standardized. To estimate the variance components of our LMM, we use a Method-of-Moments (MoM) estimator that searches for parameter values so that the population moments are close to the sample moments. Since E [***y***] = 0, we derived the MoM estimates by equating the population covariance to the empirical covariance. The population covariance is given by:

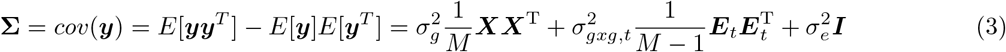

Using ***yy***^T^ as our estimate of the empirical covariance, we need to solve the following least squares problem to estimate the variance parameters :

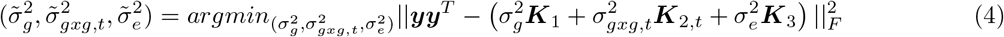

where 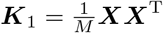 and ***K***_3_=***I***_*N*_

We show that the MoM estimator satisfies the following normal equations (see Lemma 1 in Supplementary Notes):

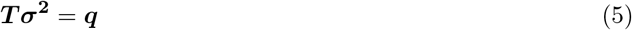

where ***T*** is a 3 *×* 3 matrix with entries *T*_*kl*_ = *tr*(***K***_*k*_***K***_*l*_), *k, l* ∈ {1, 2, 3}, *tr*() denotes the trace of the matrix, and ***q*** is a 3-vector with entries *q*_*k*_ = ***y***^*T*^ ***K***_*k*_***y***.

To compute the variance components 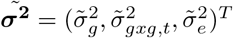, we have:

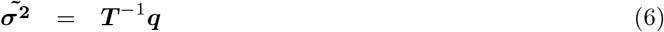

Beyond the point estimates, we also need to compute confidence intervals for 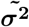 which, in turn, allow us to test the hypothesis of no ME 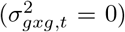. To do this, we compute the covariance matrix of 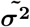 as (see Lemma 2 in Supplementary Notes):

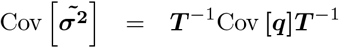

where

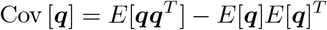

such that

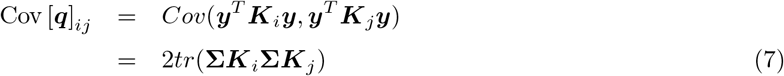

### 4.2 Efficient computation of variance components

Computing the coefficients *tr*(***K***_*k*_***K***_*l*_) of the system of linear equation 5 and Cov [***q***]_*ij*_ require 𝒪 (*N* ^2^*M*) time complexity and 𝒪 (*NM*) memory usage imposing challenging memory and computation requirements for Biobank-scale data (*N* in the hundreds of thousands, *M* in the millions). To test hypotheses, we need to compute p-values. This requires computing the point estimate and standard error which is appropriate when we have large sample sizes *N* .

To obtain an efficient estimate of 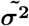, we approximate each of the coefficients of the matrix ***T*** which involves computing the trace of a matrix by an unbiased trace estimator [59]. Specifically, we estimate *T*_*i,j*_ as follows:

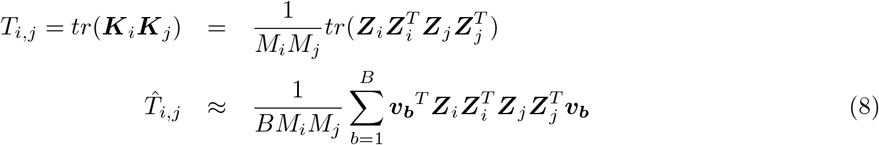

where each ***v***_***b***_ is an independent random vector with mean zero and covariance ***I***_*N*_, *B* is the total number of random vectors used for the approximation, and ***Z***_*i*_ = ***X*** or ***E*** with *M*_*i*_ columns.

To estimate Cov [***q***]_*ij*_ efficiently, we replace **Σ** in Equation 7 to obtain the plug-in estimate of Cov [***q***]_*kl*_:

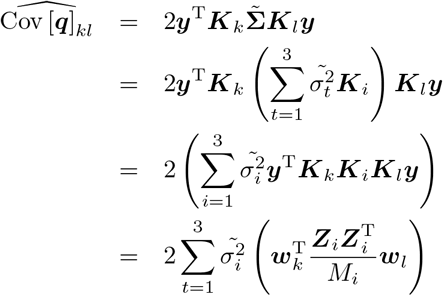

where ***w***_*i*_ = ***K***_*i*_***y***, *i* ∈ {1, …, 3}.

The computation of 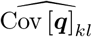 can be performed in 𝒪 (*NM*) time. Multiplication of ***w***_*k*_ = ***K***_*k*_***y*** can be decomposed into 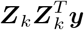 which can be computed in 𝒪 (*MN*) time. Further, by leveraging the fact that the matrices are discrete-valued genotype matrices, we can improve the time complexity of matrix-vector multiplication from 𝒪 (*NM*) to 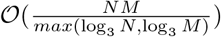 by using the Mailman algorithm [60] . Hence 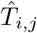 and 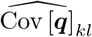 can be computed in time 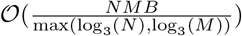 and 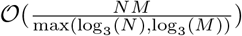 respectively. FAME uses a streaming implementation that does not require all the genotypes to be stored in memory leading to scalable memory requirements with 𝒪 (*NS*) where *S* is the number of SNPs per each stream block . We have shown that we can compute point estimates and the corresponding standard errors in sub-linear time with respect to sample size *N* and number of SNPs *M* . Therefore, we can apply our method to data sets with high sample sizes and test for the existence of ME. Finally, we note that FAME can also account for fixed-effects covariates such as age, sex, and genetic principal components (PCs) (Supplementary Information Section S2).

### 4.3 Simulations

#### Simulations to assess power and accuracy

We designed simulations to assess the power of FAME and the accuracy of its ME variance components estimates. We used the following generative model:

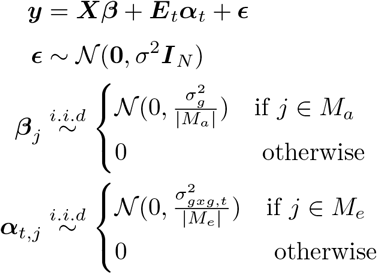

where *β*_*j*_ and *α*_*t,j*_ denotes the *j*^*th*^ element in the respective vectors of effect sizes. We set 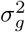 to 0.3 which is approximately the median value of the additive heritability across all the traits that we analyzed in this study. We varied the value of 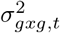 from 0.001 to 0.1. We randomly selected 10% of the SNPs to be causal for the additive effects (assigned to the indicator set *M*_*a*_) and 10% of the SNPs to be causal for the ME effect (assigned to the indicator set *M*_*e*_ which do not overlap with *M*_*a*_ and fall outside the LD block of the target SNP). As target SNP, we selected three representative SNPs with different MAF values (MAF ∈ {1%, 14%, 49%} respectively). For computational convenience, we limited our analysis to chromosomes 12 and 20 of the UKBB data, which we used as our ***X*** matrix. We simulated 1, 000 replicates for each setting. In order to assess the accuracy of the ME variance component estimates obtained by FAME, we used the same simulations as above. We then assumed that the target SNPs were known and then estimated the ME effect by partitioning the SNPs in ***X*** into two bins, LD block, which contains all the SNPs within the LD region of the target SNP; LD removed block, which contains all the SNPs outside of the LD region of the target SNP. We then used FAME to jointly fit the additive effect for both regions while only fitting the ME effect on the LD-removed region. Finally, we compared the estimated ME variance components with the ground truth.

#### Regional simulation and estimation

To localize the ME signal, we partitioned the whole genome into the region with all the SNPs lying in the same chromosome as the target SNP but outside of the LD block (termed as *local*) and all the SNPs lying on chromosomes different from the one with the target SNP (termed as *distal*). To validate the calibration of FAME when applied to test the ME effect on a specified region, we used the simulation with a total heritability of 0.25 and ratio of causal SNPs of 1%. We applied FAME to estimate the calibration of 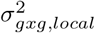 and 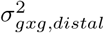 respectively.

### 4.4 Runtime comparisons

All experiments used a machine equipped with AMD EPYC 7501 32-Core Processor and a runtime budget of 3 days was provided to all tested methods.

### 4.5 Datasets

#### Simulation dataset

We obtained a set of *N* = 291, 273 unrelated white British individuals measured at *M* = 459, 792 common SNPs genotyped on the UK Biobank Axiom array to use in simulations by extracting individuals that are *>* 3rd-degree relatives and excluding individuals with putative sex chromosome aneuploidy. Unless otherwise specified, all simulations were conducted using this dataset.

#### UKBB genotypes

For analysis of real traits, we restricted our analysis to SNPs that were presented in the UK Biobank Axiom array used to genotype the UK Biobank. SNPs with greater than 1% missingness and minor allele frequency smaller than 1% were removed. Moreover, SNPs that fail the Hardy-Weinberg test at significance threshold 10^−7^ were removed. We restricted our study to self-reported British white ancestry individuals which are *>* 3^*rd*^ degree relatives that is defined as pairs of individuals with kinship coefficient *<* 1*/*2^(9*/*2)^ [61]. Furthermore, we removed individuals who are outliers for genotype heterozygosity and/or missingness and excluded SNPs that fall within the MHC region. Finally, we obtained a set of *N* = 291, 273 individuals and *M* = 454, 207 SNPs for real data analyses. We used this dataset in our analyses unless specified otherwise.

We also analyzed imputed genotypes across *N* = 291, 273 unrelated white British individuals. We removed SNPs with greater than 1% missingness, minor allele frequency smaller than 1%, SNPs that fail the Hardy-Weinberg test at significance threshold 10^−7^ as well as SNPs that lie within the MHC region (Chr6: 25–35 Mb) to obtain 4, 824, 392 SNPs.

#### Covariates and phenotypes

We selected 53 quantitative traits in the UKBB. The selected phenotypes span eight known categories: Anthropometry, Blood Biochemistry, Bone, Cardiovascular, Diabetes, Eye, Liver, and Renal. We included sex, age, and the top 20 genetic principal components (PCs) as covariates in our analysis for all phenotypes. Extra covariates were added for diastolic/systolic blood pressure (adjusted for cholesterol-lowering medication, blood pressure medication, insulin, hormone replacement therapy, and oral contraceptives). We used the PCs computed in the UKBB from a superset of 488, 295 individuals. Following prior studies, all traits were inverse rank normalized [62, 63].

## Data availability

The UK Biobank dataset used in this study is not publicly available but can be obtained by application (https://www.ukbiobank.ac.uk/).

## Code availability

FAME can be found at https://github.com/sriramlab/FAME. The simulator used in the experiments can be found at https://github.com/alipazokit/simulator. MAPIT can be found at https://github.com/lorinanthony/MAPIT.

## Acknowledgments

This research was conducted using the UK Biobank Resource under application 33127. We thank the participants of UK Biobank for making this work possible. This work was supported, in part, by NIH grants GM125055 (B.F., A.P., and S.S) and HG006399 (S.S.), and NSF grant CAREER-1943497 (B.F., A.P., and S.S.). N.Z. was supported by NIH grants R01MH130581, U01MH126798, R01MH122688, and R01GM142112.

## Notes

### Competing Interest Statement

The authors have declared no competing interest.

