## Supplementary materials for "A biobank-scale test of marginal epistasis reveals genome-wide signals of polygenic epistasis"

### Supplementary Information

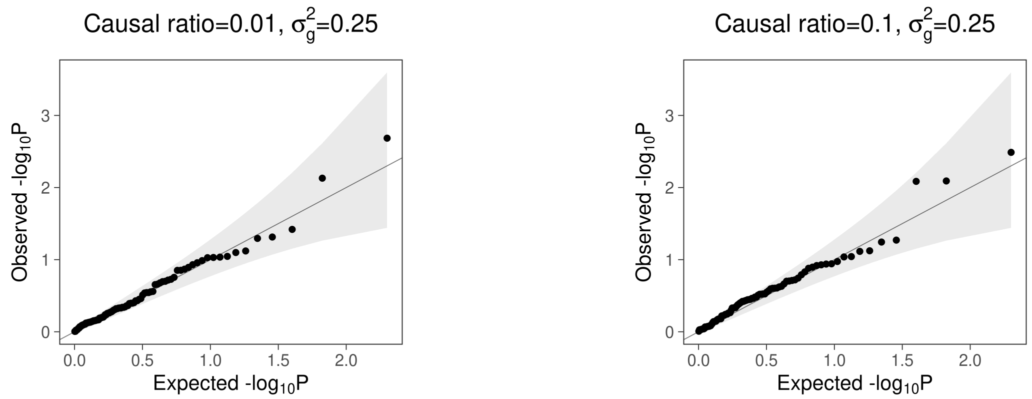


Figure S1: **Calibration of p-values for the test of ME under null simulations**. We simulated phenotypes with additive genetic effects but no genetic interactions. We applied FAME to 100 randomly selected SNPs from common SNPs on the UKBB genotyping array.

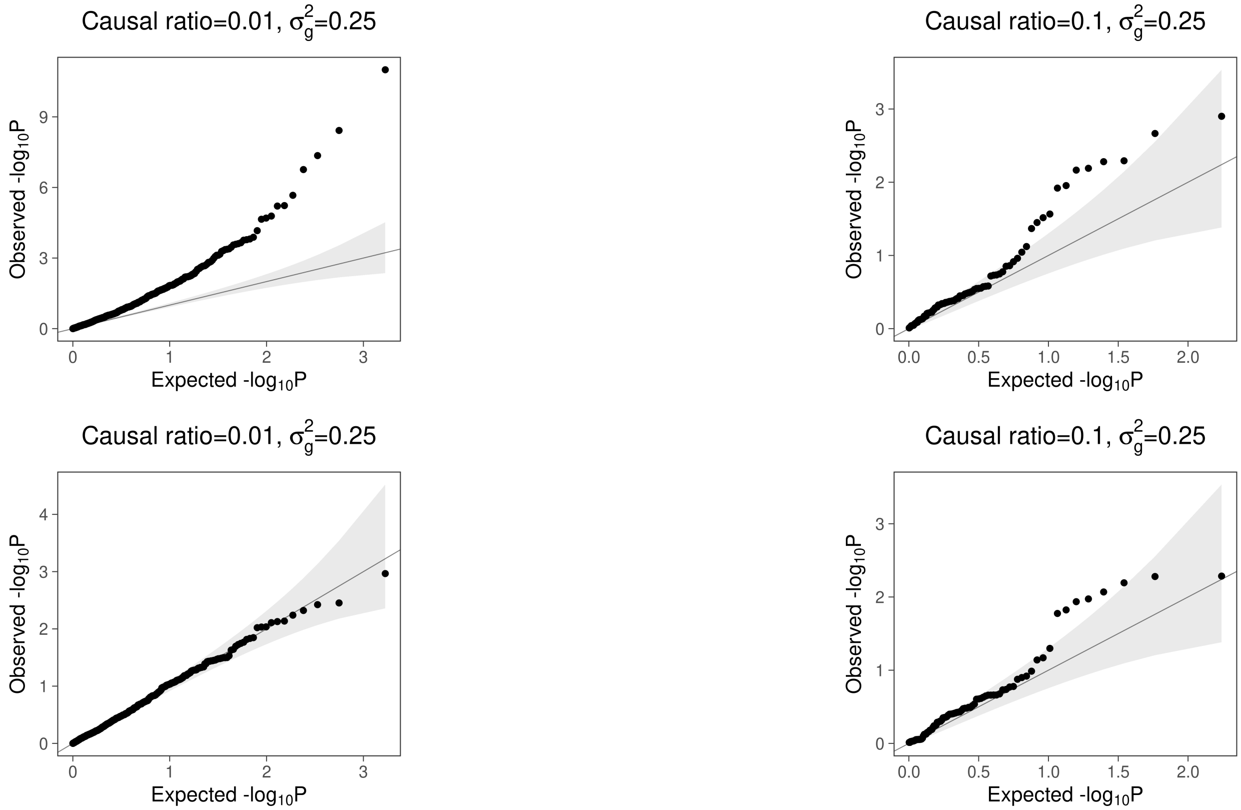


Figure S2: **Calibration of p-values for the test of ME based on inclusion of the LD block surrounding the target SNP**. In this experiment, we compared the results of applying FAME in two different scenarios. In scenario one, we constructed the ME matrix by multiplying the genotypes at each target SNP with the remaining SNPs in the genome and estimated the ME variance component and corresponding p-values (top row). In scenario two, the ME matrix is constructed by multiplying the genotype at each target SNP with all other SNPs except those within the LD block of the target SNP (bottom row). In both experiments, the additive component ($\sigma_g^2$) for all SNPs is jointly fitted with the ME matrix $\sigma^2_{gxg,t}$. The simulated phenotype has additive but no genetic interactions. The target SNPs were selected by running a GWAS.

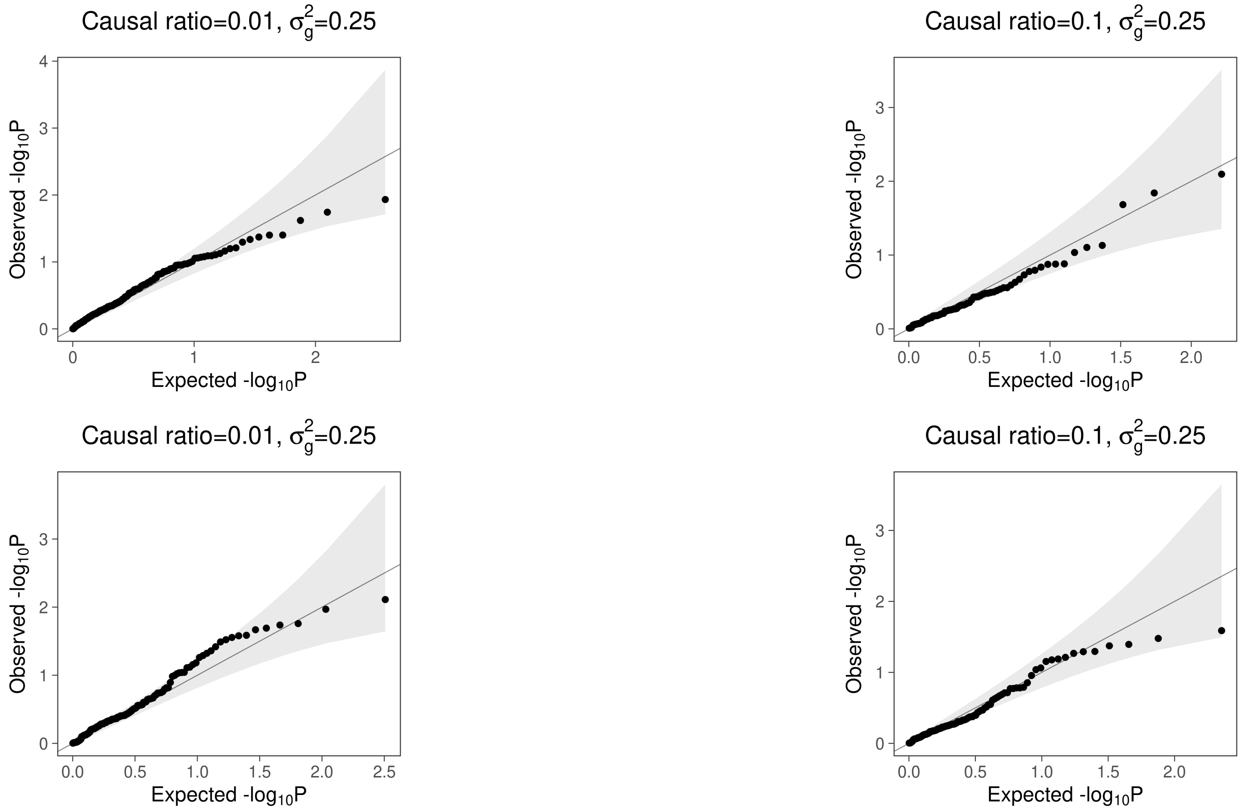


Figure S3: **Calibration of p-values in a setting where causal SNPs are imperfectly tagged**. We simulated phenotypes with different genetic architectures using imputed genotype data ($N = 291,273, M = 4,824,392$). Tests of ME were performed on array data ($N = 291,273, M = 459,792$) to model the scenario where causal SNPs are imperfectly tagged. To comprehensively understand the calibration of FAME under this scenario, we ran two simulations in each setting using different seeds (corresponding to each row).

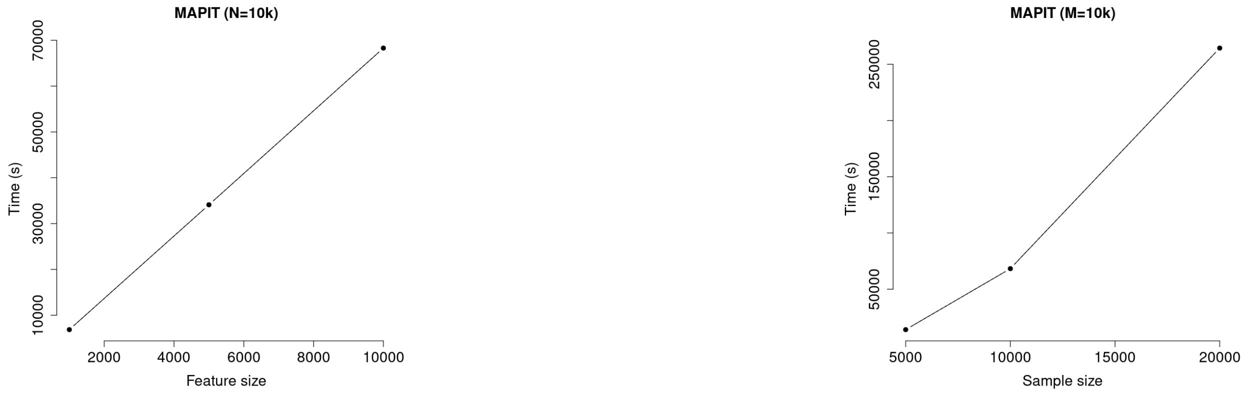


Figure S4: **Runtime for MAPIT**. We tested MAPIT on the UKBB whole-genome array data by fixing the sample size to be 10K and varying the number of SNPs (left); and by fixing the number of SNPs to 10K and varying sample size (right).

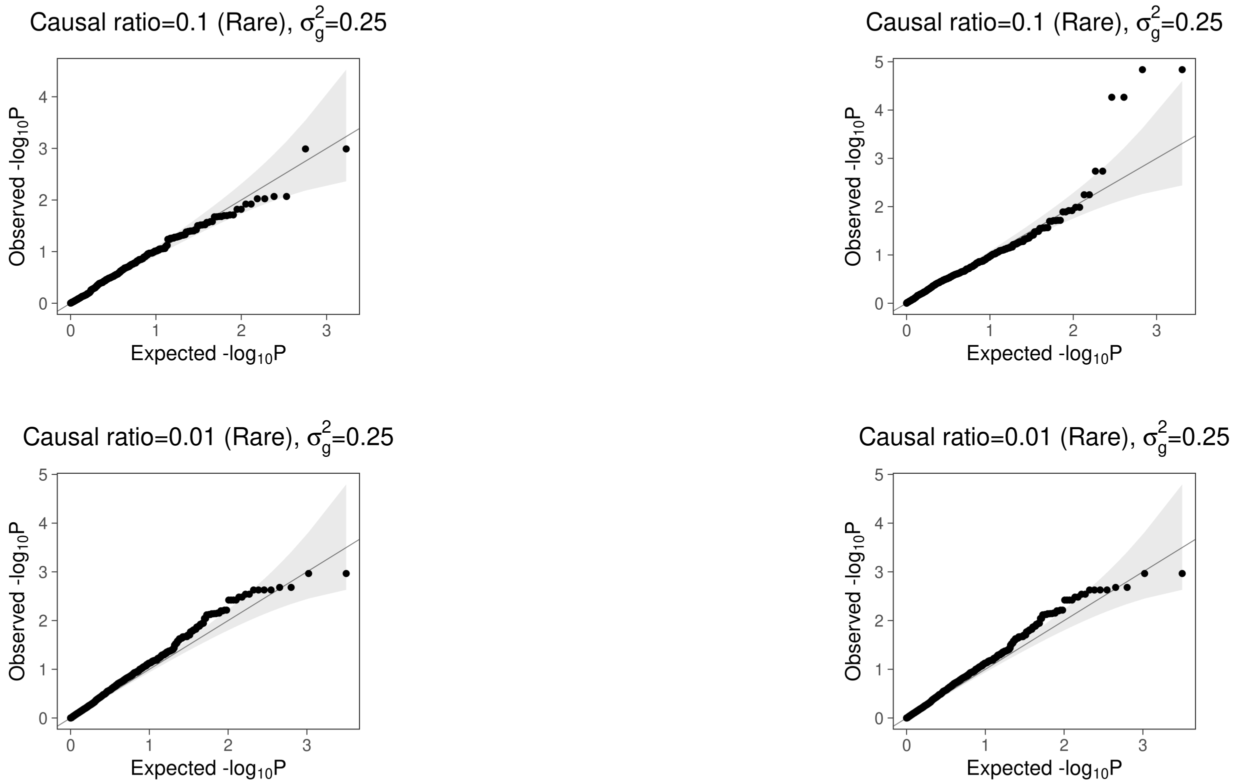


Figure S5: **Calibration of test of ME causal variants are chosen to be rare**. In this experiment, we simulated traits with no genetic interactions but additive effects (heritability= 0.25 and fraction of causal SNPs $\in \{0.1, 0.01\}$). However, we limited the causal variants to relatively rare variants (MAF $\in [0.01, 0.05]$). The MAF distribution of GWAS hits which are the target SNPs for ME has a mean of 0.16 and median of 0.09 for the Causal ratio=0.1 setting (top row); and a mean of 0.16 and median of 0.08 for the Causal ratio=0.01 setting (bottom row)

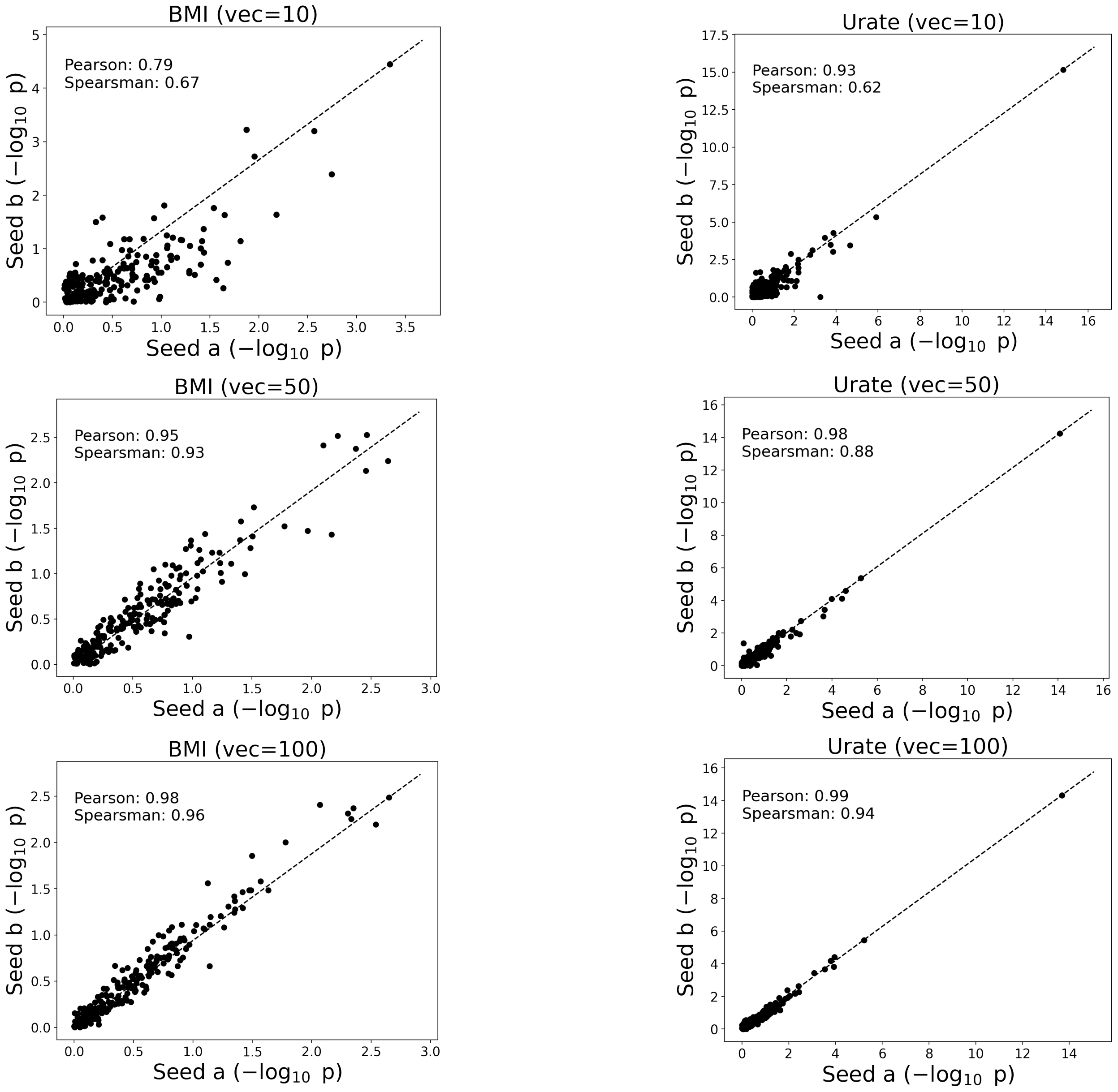


Figure S6: **Stability test of the p-values**. We selected two representative traits: BMI (no genome-wide significant ME signal) and Urate (significant ME). We reran FAME with two different sets of seeds to test the trait-SNP pairs for the above-mentioned traits. We varied the number of random vectors (vec) and found that 100 random vectors lead to highly correlated estimates across runs ($\geq 0.9$ for both traits)

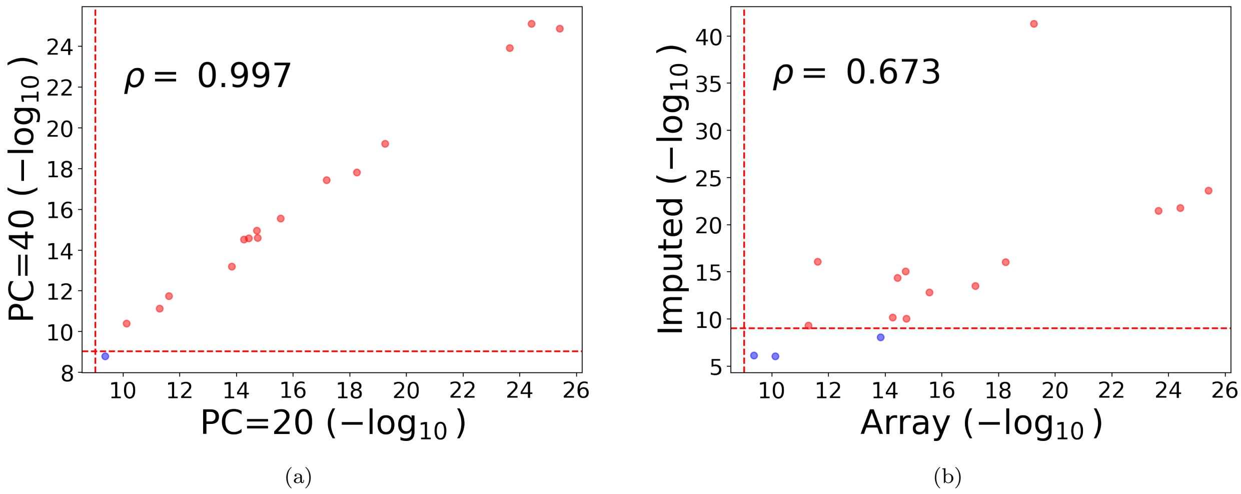


Figure S7: **Robustness tests of ME signals** (a) We assessed the robustness of ME signals to population stratification. We test the trait-loci pairs which were significant for ME signals and repeated the test by varying the number of principal components (PC=20 vs. PC=40). We plot the p-values from both analyses. (b) We assessed the robustness of ME signals to untagged SNPs in the UKBB SNP array by testing trait-loci pairs which were significant for ME in the UKBB array on imputed genotypes.

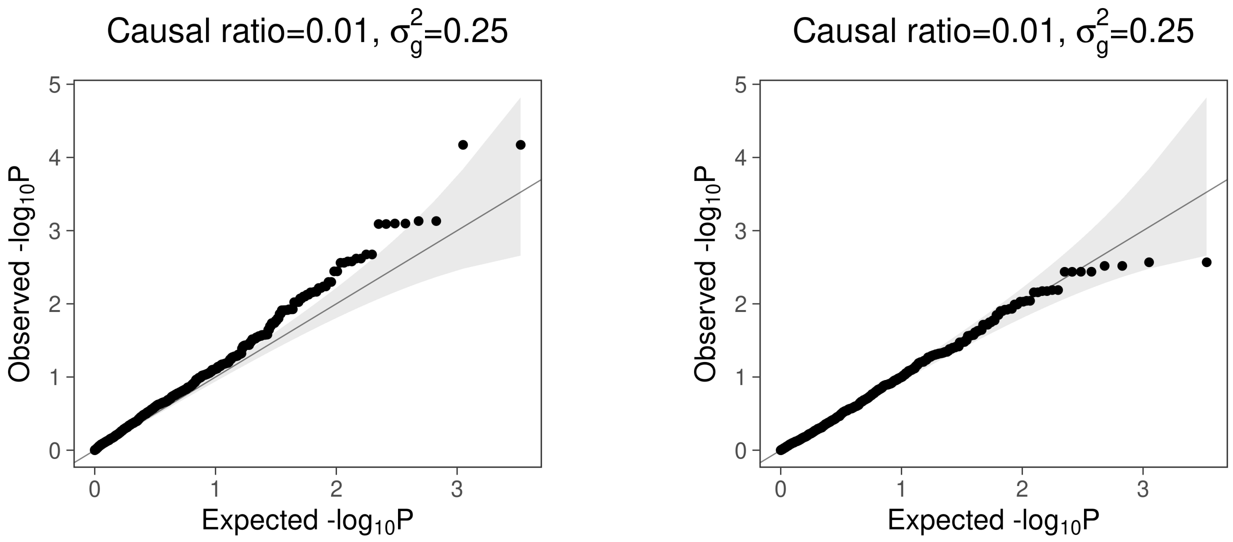


Figure S8: **QQ-plot of tests to localize ME in simulations**. We simulated phenotypes that consist only of linear additive effects based on genotypes from unrelated white-British individuals in the UKBB (fraction of causal SNPs = 0.01 and heritability= 0.25). The target SNPs were selected to be significant SNPs obtained using a GWAS. The figure on the left shows the QQ-plot of test for ME that lies on the same chromosome as the target SNP (with the LD block around the target SNP removed) (*local*). The figure on the right shows the QQ-plot of a test for ME effects that lie on a different chromosome than the target SNP (*distal*). No significant ME signals ($p \leq 5 \times 10^{-8}$) were found across all the settings.

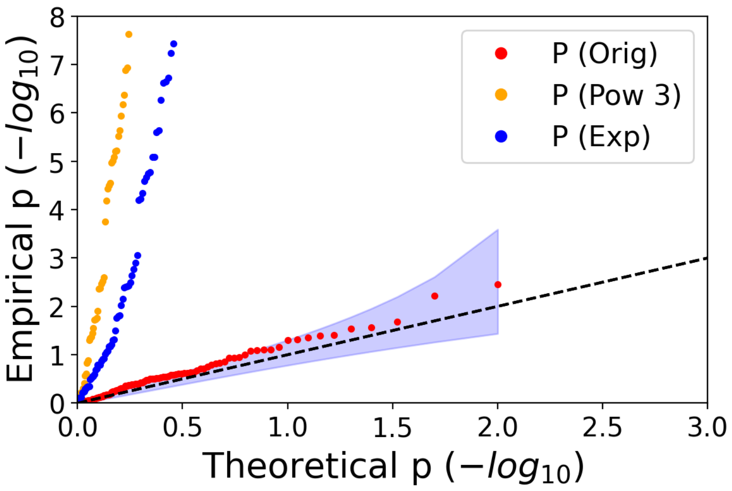


Figure S9: **Effect of scaling the phenotype on ME estimates**. We randomly selected 100 GWAS significant SNPs associated with height and tested them for ME with different scaling (Orig: original scale; Pow 3, the third power of the target trait; Exp, the exponent of the target trait).

### S1 Estimating heritability from ME variance component estimates

Here we compute the heritability explained by the ME variance component estimated in Equation 6 following the derivation in [64].

Under the assumption that all matrices are standardized, the heritability associated with ME effects at SNP $t$ can be computed as:

$$\hat{h}^2_{gxg,t} = \frac{\hat{\sigma}^2_{gxg,t}}{\hat{\sigma}^2_g + \hat{\sigma}^2_{gxg,t} + \hat{\sigma}^2_e}$$

However, if the matrix $\mathbf{E} = \mathbf{X}_{-t} \odot \mathbf{X}_t$ is not standardized, then we need to derive a more general formula.

Let $V_T$ denote the total phenotypic variance.

$$V_T = \frac{\sum_i y_i^2}{N} - \Big(\frac{\sum_i y_i}{N}\Big)^2 \tag{9}$$

We can re-write our model as follows:

$$y_i = \sum_{j=1}^{M} x_{ij}\beta_j + \sum_{k=1}^{M-1} e_{ik}\alpha_k + \epsilon_i$$

Here $e_{ik}$ is the element in the $i^{th}$ row and $k^{th}$ column of $\boldsymbol{E}$. Based on the distributional assumptions of our model, the following holds:

$$\begin{aligned}
\mathbb{E}[\beta_j\beta_{j'}] = 0 \text{ for } j \neq j', \quad \mathbb{E}[\beta_j^2] = \frac{\sigma_g^2}{M} \\
\mathbb{E}[\alpha_k\alpha_{k'}] = 0 \text{ for } k \neq k', \quad \mathbb{E}[\alpha_k^2] = \frac{\sigma^2_{gxg,t}}{M'} \\
\mathbb{E}[\epsilon_i\epsilon_{i'}] = 0 \text{ for } i \neq i', \quad \mathbb{E}[\epsilon_i^2] = \sigma^2_\epsilon \\
\mathbb{E}[\beta_j\alpha_k] = 0; \quad \mathbb{E}[\beta_j\epsilon_i] = 0; \quad \mathbb{E}[\alpha_k\epsilon_i] = 0;
\end{aligned}$$

where $M' = M - 1$.

To compute the expectation of the terms in the first summation of Equation 9:

$$\begin{aligned}
\mathbb{E}[y_i^2] &= \mathbb{E}[(\sum_{j=1}^{M} x_{ij}\beta_j + \sum_{k=1}^{M'} e_{ik}\alpha_k + \epsilon_i)^2] \\
&= \sum_{j=1}^{M} x_{ij}^2\mathbb{E}[\beta_{ij}^2] + \sum_{k=1}^{M'} e_{ik}^2\mathbb{E}[\alpha_k^2] + \mathbb{E}[\epsilon_i^2] \\
&= (\sum_{j=1}^{M} x_{ij}^2)\frac{\sigma_g^2}{M} + (\sum_{k=1}^{M'} e_{ik}^2)\frac{\sigma^2_{gxg,t}}{M'} + \sigma^2_\epsilon
\end{aligned}$$

This gives us:

$$\frac{1}{N}\sum_i \mathbb{E}[y_i^2] = \frac{1}{N}(\sum_{i=1}^{N}\sum_{j=1}^{M} x_{ij}^2)\frac{\sigma_g^2}{M} + \frac{1}{N}(\sum_{i=1}^{N}\sum_{k=1}^{M'} e_{ik}^2)\frac{\sigma_{gxg,t}^2}{M'} + \sigma_\epsilon^2$$
$$= \frac{tr(\boldsymbol{X}\boldsymbol{X}^T)}{N}\frac{\sigma_g^2}{M} + \frac{tr(\boldsymbol{E}\boldsymbol{E}^T)}{N}\frac{\sigma_{gxg,t}^2}{M'} + \sigma_\epsilon^2 \quad (10)$$

The expected value for the second summation in Equation 9 is:

$$\mathbb{E}[(y_1 + ... + y_N)^2] \quad (11)$$
$$= \mathbb{E}[\sum_{ii'} x_{i1}x_{i'1}\beta_1^2 + ... + \sum_{ii'} x_{iM}x_{i'M}\beta_M^2 + \sum_{ii'} e_{i1}e_{i'1}\alpha_1^2 + ... + \sum_{ii'} e_{iM'}e_{i'M'}\alpha_{M'}^2 + \sum_i \epsilon_i^2]$$
$$= (\sum_j\sum_{ii'} x_{ij}x_{i'j})\frac{\sigma_g^2}{M} + (\sum_k\sum_{ii'} e_{ik}e_{i'k})\frac{\sigma_{gxg,t}^2}{M'} + N\sigma_\epsilon^2$$
$$= sum(\boldsymbol{X}\boldsymbol{X}^T)\frac{\sigma_g^2}{M} + sum(\boldsymbol{E}\boldsymbol{E}^T)\frac{\sigma_{gxg,t}^2}{M'} + N\sigma_\epsilon^2 \quad (12)$$

Therefore

$$\mathbb{E}[V_T] = \frac{tr(\boldsymbol{X}\boldsymbol{X}^T)}{N}\frac{\sigma_g^2}{M} + \frac{tr(\boldsymbol{E}\boldsymbol{E}^T)}{N}\frac{\sigma_{gxg,t}^2}{M'} + \sigma_\epsilon^2 - \frac{sum(\boldsymbol{X}\boldsymbol{X}^T)}{N^2}\frac{\sigma_g^2}{M} - \frac{sum(\boldsymbol{E}\boldsymbol{E}^T)}{N^2}\frac{\sigma_{gxg,t}^2}{M'} - \frac{1}{N}\sigma_\epsilon^2$$
$$= \sigma_g^2 + \left(\frac{tr(\boldsymbol{E}\boldsymbol{E}^T)}{N} - \frac{sum(\boldsymbol{E}\boldsymbol{E}^T)}{N^2}\right)\frac{\sigma_{gxg,t}^2}{M'} + \frac{N-1}{N}\sigma_\epsilon^2$$

where the last equation holds under the assumption that $\boldsymbol{X}$ is standardized.

We define the residual with respect to the ME variance component as $r_i = \sum_{j=1}^{M} x_{ij}\beta_j + \epsilon_i$ so that the residual variance is:

$$\mathbb{E}[V_R] = \frac{tr(\boldsymbol{X}\boldsymbol{X}^T)}{N}\frac{\sigma_g^2}{M} + \sigma_\epsilon^2 - \frac{sum(\boldsymbol{X}\boldsymbol{X}^T)}{N^2}\frac{\sigma_g^2}{M} - \frac{1}{N}\sigma_\epsilon^2$$
$$= \sigma_g^2 + \frac{N-1}{N}\sigma_\epsilon^2$$

Finally, the expected ME heritability can be computed as

$$\mathbb{E}[h_{gxg,t}^2] \approx 1 - \frac{\mathbb{E}[V_R]}{\mathbb{E}[V_T]}$$
$$= 1 - \frac{\sigma_g^2 + \frac{N-1}{N}\sigma_\epsilon^2}{\sigma_g^2 + (\frac{tr(\boldsymbol{E}\boldsymbol{E}^T)}{N} - \frac{sum(\boldsymbol{E}\boldsymbol{E}^T)}{N^2})\frac{\sigma_{gxg,t}^2}{M'} + \frac{N-1}{N}\sigma_\epsilon^2}$$
$$= \frac{(\frac{tr(\boldsymbol{E}\boldsymbol{E}^T)}{N} - \frac{sum(\boldsymbol{E}\boldsymbol{E}^T)}{N^2})\frac{\sigma_{gxg,t}^2}{M'}}{\sigma_g^2 + (\frac{tr(\boldsymbol{E}\boldsymbol{E}^T)}{N} - \frac{sum(\boldsymbol{E}\boldsymbol{E}^T)}{N^2})\frac{\sigma_{gxg,t}^2}{M'} + \frac{N-1}{N}\sigma_\epsilon^2}$$
$$= \frac{\mathcal{C}\sigma_{gxg,t}^2}{\sigma_g^2 + \mathcal{C}\sigma_{gxg,t}^2 + \frac{N-1}{N}\sigma_\epsilon^2}$$

where $\mathcal{C} \equiv (\frac{tr(\boldsymbol{E}\boldsymbol{E}^T)}{N} - \frac{sum(\boldsymbol{E}\boldsymbol{E}^T)}{N^2})\frac{1}{M'}$.

We plug-in the estimates $\tilde{\boldsymbol{\sigma}}^2$ to obtain

$$\tilde{h}^2_{gxg} = \frac{\tilde{\mathcal{C}}\tilde{\sigma}^2_{gxg}}{\tilde{\sigma}^2_g + \tilde{\mathcal{C}}\tilde{\sigma}^2_{gxg} + \frac{N-1}{N}\tilde{\sigma}^2_\epsilon}$$

where $\tilde{\mathcal{C}}$ is computed efficiently using the randomized trace estimates.
During the actual computation, $tr(\boldsymbol{EE}^T)$ can be directly accessed via the model intermediate output while we compute $sum(\boldsymbol{EE}^T)$ as:

$$\begin{aligned}
sum(\boldsymbol{EE}^T) &= \sum_{ii'}(\boldsymbol{EE}^T)_{ii'} \\
&= \sum_{ii'}\sum_{j} e_{ij}e_{i'j} \\
&= \sum_{j}\sum_{ii'} e_{ij}e_{i'j} \\
&= \sum_{j}(\sum_{i} e_{ij})(\sum_{i'} e_{i'j}) \\
&= \sum_{j}(\sum_{i} e_{ij})^2 \\
&= N^2\sum_{j} corr(\boldsymbol{X}_{:t}, \boldsymbol{X}_{:j})^2
\end{aligned}$$

where the last equation holds because

$$\begin{aligned}
e_{ij} &= x_{it}x_{ij} \\
\sum_i e_{ij} &= \sum_i x_{it}x_{ij} \\
&= NCorr(\boldsymbol{X}_{:t}, \boldsymbol{X}_{:j}) \qquad (13)
\end{aligned}$$

and Equation 13 holds under the assumption that $\mathbf{X}$ is column-wise standardized. In practice, we estimate the squared correlation with Plink [65].

To further estimate the standard error of $h^2_{gxg}$, we use the fact that the target traits have been standardized; therefore, $V_T \approx 1$, and thus

$$\begin{aligned}
Var(h^2_{gxg}) &= Var(1 - \frac{\mathbb{E}[V_R]}{\mathbb{E}[V_T]}) \\
&\approx Var(1 - \mathbb{E}[V_R]) \\
&= Var(\mathbb{E}[V_R]) \\
&= Var(\mathcal{C}\sigma^2_{gxg}) \\
&= \mathcal{C}^2 Var(\sigma^2_{gxg})
\end{aligned}$$

Therefore the corresponding estimation for standard error $\widetilde{SE}(h^2_{gxg}) = \mathcal{C} * \widetilde{SE}(\sigma^2_{gxg})$.

#### S2 Including covariates

We can extend each of our models to include covariates as follows:

$$\boldsymbol{y} = \boldsymbol{W}\boldsymbol{\alpha} + \sum_k \boldsymbol{Z}_k\boldsymbol{\beta}_k + \boldsymbol{\epsilon} \tag{14}$$

Here $\boldsymbol{W}$ is a $N \times C$ matrix of covariates while $\boldsymbol{\alpha}$ is a vector of fixed effects of length $C$. Matrix $\boldsymbol{Z}_k$ can represent genotype or GxE matrices. In this setting, we need to solve the following normal equations to estimate the variance components.

$$\begin{bmatrix} \boldsymbol{T} & \boldsymbol{b} \\ \boldsymbol{b}^T & N - C \end{bmatrix} \begin{bmatrix} \sigma_1^2 \\ \vdots \\ \sigma_k^2 \\ \sigma_e^2 \end{bmatrix} = \begin{bmatrix} \boldsymbol{c} \\ \boldsymbol{y}^T\boldsymbol{V}\boldsymbol{y} \end{bmatrix} \tag{15}$$

Here $\boldsymbol{V} = \boldsymbol{I}_N - \boldsymbol{W}(\boldsymbol{W}^T\boldsymbol{W})^{-1}\boldsymbol{W}^T$ and $\boldsymbol{T}$ is a $K \times K$ matrix where $T_{k,l} = tr(\boldsymbol{K}_k\boldsymbol{V}\boldsymbol{K}_l\boldsymbol{V})$, and $b$ is a vector of length $K$ where $b_k = tr(\boldsymbol{V}\boldsymbol{K}_k)$, and $\boldsymbol{c}$ is a vector of length $K$ where $c_k = \boldsymbol{y}^T\boldsymbol{V}\boldsymbol{K}_k\boldsymbol{V}\boldsymbol{y}$, $\boldsymbol{K}_k = \frac{\boldsymbol{Z}_k\boldsymbol{Z}_k^T}{M}$ where $M$ is the number of column in $Z_k$. Commonly, the number of covariates $C$ is small (tens to hundreds) so that including covariates does not significantly affect the computational cost. The cost of computing the elements of the normal equations 15 includes the cost of inverting $\boldsymbol{W}^T\boldsymbol{W}$ which is a $C \times C$ matrix and multiplying $\boldsymbol{W}$ by a real-valued vector of length $N$ can be computed in $\mathcal{O}(C^3 + NC)$.

### S3 Proofs

**Lemma 1.**

$$(\tilde{\sigma}_g^2, \tilde{\sigma}_{gxg,t}^2, \tilde{\sigma}_e^2) = argmin_{(\sigma_g^2, \sigma_{gxg,t}^2, \sigma_e^2)} ||\boldsymbol{y}\boldsymbol{y}^T - \left(\sigma_g^2 \boldsymbol{K}_1 + \sigma_{gxg,t}^2 \boldsymbol{K}_{2,t} + \sigma_e^2 \boldsymbol{K}_3\right)||_F^2 \tag{16}$$

*satisfies the following normal equations:*

$$\boldsymbol{T}\boldsymbol{\sigma^2} = \boldsymbol{q} \tag{17}$$

*where* $\boldsymbol{K}_1 = \frac{1}{M}\boldsymbol{X}\boldsymbol{X}^T$*,* $\boldsymbol{K}_{2,t} = \frac{1}{M-1}\boldsymbol{E}_t\boldsymbol{E}_t^T$ *and* $\boldsymbol{K}_3 = \boldsymbol{I}_N$*,* $\boldsymbol{T}$ *is a* $3 \times 3$ *matrix with entries* $T_{kl} = tr(\boldsymbol{K}_k\boldsymbol{K}_l), k, l \in \{1, 2, 3\}$*,* $tr()$ *denotes the trace of the matrix, and* $\boldsymbol{q}$ *is a* 3*-vector with entries* $c_k = \boldsymbol{y}^T\boldsymbol{K}_k\boldsymbol{y}$*.*

*Proof.* Using the definition of the Frobenius norm of a matrix $\boldsymbol{A}$ ($||\boldsymbol{A}||_F = \sqrt{tr[\boldsymbol{A}\boldsymbol{A}^T]}$), the above equation can be re-written as:

$$(\tilde{\sigma}_g^2, \tilde{\sigma}_{gxg,t}^2, \tilde{\sigma}_e^2) = argmin_{(\sigma_g^2, \sigma_{gxg,t}^2, \sigma_e^2)} tr[(\boldsymbol{y}\boldsymbol{y}^T - \left(\sigma_g^2\boldsymbol{K}_1 + \sigma_{gxg,t}^2\boldsymbol{K}_{2,t} + \sigma_e^2\boldsymbol{K}_3\right))(\boldsymbol{y}\boldsymbol{y}^T - \left(\sigma_g^2\boldsymbol{K}_1 + \sigma_{gxg,t}^2\boldsymbol{K}_{2,t} + \sigma_e^2\boldsymbol{K}_3\right))^T]$$

Let $\boldsymbol{\theta} := [\sigma_g^2, \sigma_{gxg,t}^2, \sigma_e^2]$. Then

$$\partial f(\boldsymbol{\theta})/\partial\sigma_g^2 = 0$$

$$\Rightarrow \sigma_g^2 tr(\boldsymbol{K}_1^2) + \sigma_{gxg}^2 tr(\boldsymbol{K}_1\boldsymbol{K}_{2,t}) + \sigma_e^2 tr(\boldsymbol{K}_1\boldsymbol{K}_3) = \boldsymbol{y}^T\boldsymbol{K}_1\boldsymbol{y} \tag{18}$$

Thus, the minimizer of $f(\boldsymbol{\theta})$ must satisfy Equation 18:

$$\tilde{\sigma}_g^2 tr(\boldsymbol{K}_1^2) + \tilde{\sigma}_{gxg}^2 tr(\boldsymbol{K}_1\boldsymbol{K}_{2,t}) + \tilde{\sigma}_e^2 tr(\boldsymbol{K}_1\boldsymbol{K}_3) = \boldsymbol{y}^T\boldsymbol{K}_1\boldsymbol{y} \tag{19}$$

Similarly, we can take the partial derivative of $\sigma_{gxg}^2$ and $\sigma_e^2$ to obtain:

$$\tilde{\sigma}_g^2 tr(\boldsymbol{K}_1\boldsymbol{K}_{2,t}) + \tilde{\sigma}_{gxg}^2 tr(\boldsymbol{K}_{2,t}^2) + \tilde{\sigma}_e^2 tr(\boldsymbol{K}_{2,t}\boldsymbol{K}_3) = \boldsymbol{y}^T\boldsymbol{K}_{2,t}\boldsymbol{y} \tag{20}$$

$$\tilde{\sigma}_g^2 tr(\boldsymbol{K}_1\boldsymbol{K}_3) + \tilde{\sigma}_{gxg}^2 tr(\boldsymbol{K}_{2,t}\boldsymbol{K}_3) + \tilde{\sigma}_e^2 tr(\boldsymbol{K}_3^2) = \boldsymbol{y}^T\boldsymbol{K}_3\boldsymbol{y} \tag{21}$$

For simplicity, we can compactly rewrite the above system of equations as:

$$\boldsymbol{T}\boldsymbol{\sigma^2} = \boldsymbol{q} \tag{22}$$

where $\boldsymbol{T}$ is a $3 \times 3$ matrix with entries $T_{kl} = tr(\boldsymbol{K}_k\boldsymbol{K}_l), k, l \in \{1, 2, 3\}$, $tr()$ denotes the trace of the matrix, and $\boldsymbol{q}$ is a 3-vector with entries $c_k = \boldsymbol{y}^T\boldsymbol{K}_k\boldsymbol{y}$. □

**Lemma 2.** *Let* $\boldsymbol{z} \sim \mathcal{N}(\mathbf{0}, \boldsymbol{C})$*.* $Cov\left[\boldsymbol{z}^T\boldsymbol{A}\boldsymbol{z}, \boldsymbol{z}^T\boldsymbol{B}\boldsymbol{z}\right] = 2tr(\boldsymbol{C}\boldsymbol{A}\boldsymbol{C}\boldsymbol{B})$ *for symmetric matrices* $\boldsymbol{A}$*,* $\boldsymbol{B}$*.*

*Proof.*

$$\text{Cov}\left[\boldsymbol{z}^{\mathrm{T}}\boldsymbol{A}\boldsymbol{z}, \boldsymbol{z}^{\mathrm{T}}\boldsymbol{B}\boldsymbol{z}\right] = \mathbb{E}\left[\boldsymbol{z}^{\mathrm{T}}\boldsymbol{A}\boldsymbol{z}\boldsymbol{z}^{\mathrm{T}}\boldsymbol{B}\boldsymbol{z}\right] - \mathbb{E}\left[\boldsymbol{z}^{\mathrm{T}}\boldsymbol{A}\boldsymbol{z}\right]\mathbb{E}\left[\boldsymbol{z}^{\mathrm{T}}\boldsymbol{B}\boldsymbol{z}\right] \tag{23}$$

$$\begin{aligned}
\mathbb{E}\left[\boldsymbol{z}^{\mathrm{T}}\boldsymbol{A}\boldsymbol{z}\boldsymbol{z}^{\mathrm{T}}\boldsymbol{B}\boldsymbol{z}\right] &= \mathbb{E}\left[\left(\sum_{i,j} z_i A_{ij} z_j\right)\left(\sum_{k,l} z_k B_{kl} z_l\right)\right] \\
&= \mathbb{E}\left[\sum_{i,j,k,l} A_{ij} B_{kl} z_i z_j z_k z_l\right] \\
&= \sum_{i,j,k,l} A_{ij} B_{kl} \mathbb{E}\left[z_i z_j z_k z_l\right] && (24) \\
&= \sum_{i,j,k,l} A_{ij} B_{kl} [\mathbb{E}\left[z_i z_j\right]\mathbb{E}\left[z_k z_l\right] + \mathbb{E}\left[z_i z_k\right]\mathbb{E}\left[z_j z_l\right] + \mathbb{E}\left[z_i z_l\right]\mathbb{E}\left[z_k z_j\right]] && (25) \\
&= \sum_{i,j,k,l} A_{ij} B_{kl} [C_{ij} C_{kl} + C_{ik} C_{jl} + C_{il} C_{jk}] && (26) \\
&= \sum_{i,j,k,l} A_{ij} B_{kl} C_{ij} C_{kl} + \sum_{i,j,k,l} A_{ij} B_{kl} C_{ik} C_{jl} + \sum_{i,j,k,l} A_{ij} B_{kl} C_{il} C_{jk} \\
&= \sum_{i,j,k,l} A_{ij} C_{ij} B_{kl} C_{kl} + \sum_{i,j,k,l} A_{ij} C_{ik} B_{kl} C_{jl} + \sum_{i,j,k,l} A_{ij} C_{il} B_{kl} C_{jk} \\
&= \sum_{i,j} A_{ij} C_{ij} \sum_{k,l} B_{kl} C_{kl} + \sum_{i,j,k,l} A_{ij} C_{ik} B_{kl} C_{jl} + \sum_{i,j,k,l} A_{ij} C_{ik} B_{lk} C_{jl} && (27) \\
&= \left(\sum_{i,j} A_{ij} C_{ij}\right)\left(\sum_{k,l} B_{kl} C_{kl}\right) + \sum_{i,j,k,l} A_{ij} C_{ik} B_{kl} C_{jl} + \sum_{i,j,k,l} A_{ij} C_{ik} B_{kl} C_{jl} && (28) \\
&= tr(\boldsymbol{A}\boldsymbol{C}) tr(\boldsymbol{B}\boldsymbol{C}) + 2\sum_{i,j,k,l} A_{ij} C_{ik} B_{kl} C_{jl} && (29) \\
&= tr(\boldsymbol{A}\boldsymbol{C}) tr(\boldsymbol{B}\boldsymbol{C}) + 2\sum_{j,k}\left(\sum_{i} A_{ji} C_{ik}\right)\left(\sum_{l} B_{kl} C_{lj}\right) && (30) \\
&= tr(\boldsymbol{A}\boldsymbol{C}) tr(\boldsymbol{B}\boldsymbol{C}) + 2\sum_{j,k} (\boldsymbol{A}\boldsymbol{C})_{jk} (\boldsymbol{B}\boldsymbol{C})_{kj} && (31) \\
&= tr(\boldsymbol{A}\boldsymbol{C}) tr(\boldsymbol{B}\boldsymbol{C}) + 2tr(\boldsymbol{A}\boldsymbol{C}\boldsymbol{B}\boldsymbol{C}) && (32)
\end{aligned}$$

Equation 24 follows by linearity of expectation while Equation 25 follows from an application of Isserlis' theorem and Equation 26 follows from the fact that $\boldsymbol{z} \sim \mathcal{N}(\mathbf{0}, \boldsymbol{C})$. Equation 27 follows by grouping factors in the first term and by interchanging indices $l$ and $k$ in the second term. Equation 28 follows from the fact that $\boldsymbol{B}$ is symmetric so that $B_{kl} = B_{lk}$. Equation 29 follows from the observation that the last two terms are identical. Equation 30 uses the fact that matrices $\boldsymbol{A}$ and $\boldsymbol{C}$ are symmetric. Equation 31 is the definition of matrix multiplication and Equation 32 follows from the definition of the trace.

$$\begin{aligned}
\mathbb{E}\left[\boldsymbol{z}^{\mathrm{T}}\boldsymbol{A}\boldsymbol{z}\right] &= \mathbb{E}\left[\sum_{i,j} z_i A_{ij} z_j\right] \\
&= \mathbb{E}\left[\sum_{i,j} A_{ij} z_i z_j\right] \\
&= \sum_{i,j} A_{ij}\mathbb{E}\left[z_i z_j\right] \\
&= \sum_{i,j} A_{ij} C_{ij} \\
&= tr(\boldsymbol{AC}) \qquad (33)
\end{aligned}$$

Equation 23 follows by combining Equations 32 and 33. □

Table S1: Test of robustness of p-values across other five different seeds. For all the trait-loci pairs with significant ME detected, only one trait-locus failed to pass a significant threshold across all the seeds (bold).

| Trait | CHR | SNP | $-\log_{10} P_1$ | $-\log_{10} P_2$ | $-\log_{10} P_3$ | $-\log_{10} P_4$ | $-\log_{10} P_5$ |
|---|---|---|---|---|---|---|---|
| Alanine aminotransferase | 22 | 44388817 | 14.400 | 14.890 | 14.928 | 14.716 | 14.630 |
| Apolipoprotein B | 11 | 116648917 | 10.050 | 11.057 | 10.778 | 10.216 | 10.875 |
| C-reactive protein | 1 | 66257838 | 9.720 | 9.196 | 9.274 | 9.625 | **8.809** |
| Cholesterol | 11 | 116648917 | 13.120 | 12.843 | 13.380 | 13.558 | 13.555 |
| Hemoglobin A1c | 8 | 41542093 | 14.909 | 14.480 | 14.779 | 14.881 | 14.852 |
| Lipoprotein-A | 6 | 160578069 | 19.693 | 19.704 | 19.693 | 19.693 | 19.810 |
| | | 160560845 | 18.103 | 17.140 | 17.611 | 18.103 | 17.380 |
| | 19 | 45414399 | 15.018 | 14.141 | 14.218 | 15.030 | 15.368 |
| Mean platelet volume | 20 | 57597970 | 17.858 | 18.258 | 17.487 | 18.026 | 18.352 |
| Monocyte count | 13 | 28623048 | 23.596 | 23.614 | 22.795 | 22.882 | 23.373 |
| SHBG | 17 | 7145117 | 24.285 | 24.863 | 25.601 | 25.226 | 25.307 |
| | | 7254315 | 15.608 | 15.529 | 15.529 | 15.919 | 15.759 |
| Testosterone | 7 | 99032593 | 11.727 | 11.296 | 11.326 | 11.181 | 11.792 |
| | 12 | 2977954 | 25.748 | 25.471 | 25.915 | 24.867 | 26.183 |
| Triglycerides | 11 | 116648917 | 12.209 | 11.708 | 11.883 | 11.938 | 12.360 |
| Urate | 1 | 145630111 | 14.767 | 14.385 | 14.615 | 14.493 | 14.587 |

Table S2: Comparison of average MAF at significant ME and GWAS loci for each trait. $g$ count is the number of GWAS-significant SNPs ($p \leq 5 \times 10^{-8}$.); Among these trait-loci pairs, $gxg$ Count is the number of SNPs significant for ME ($p \leq \frac{5\times10^{-8}}{53}$). Average MAF ($g$) is the average minor allele frequency at GWAS-significant SNPs; Average MAF ($gxg$) is the average minor allele frequency at the significant ME SNPs. The average MAF ($gxg$) across all trait-loci pairs (a total of 16) is 0.11; the average MAF ($g$) across all trait-loci pairs (a total of $15,601$) is 0.24.

| Trait | $g$ Count | $gxg$ Count | Average MAF ($g$) | Average MAF ($gxg$) |
|---|---|---|---|---|
| Alanine aminotransferase | 152 | 1 | 0.26 | 0.18 |
| Apolipoprotein B | 237 | 1 | 0.21 | 0.13 |
| C-reactive protein | 253 | 1 | 0.22 | 0.20 |
| Cholesterol | 237 | 1 | 0.23 | 0.13 |
| Hemoglobin A1c | 444 | 1 | 0.25 | 0.03 |
| Lipoprotein-A | 104 | 3 | 0.14 | 0.18 |
| Mean platelet volume | 898 | 1 | 0.22 | 0.05 |
| Monocyte count | 458 | 1 | 0.24 | 0.03 |
| SHBG | 323 | 2 | 0.23 | 0.07 |
| Testosterone | 84 | 2 | 0.22 | 0.06 |
| Triglycerides | 294 | 1 | 0.22 | 0.13 |
| Urate | 269 | 1 | 0.22 | 0.01 |

Table S3: Localizing marginal epistasis. *local* denotes ME effects of the target SNP paired with SNPs that lie on the same chromosome while *distal* denotes ME effects of the target SNP paired with SNPs in different chromosomes. Entries with bold text indicates the corresponding ME estimates were statistically significant ($p \leq \frac{5\times10^{-8}}{53}$)

| Trait | CHR | SNP | $gxg$ $\times 0.001$ | $gxg_{local}$ $\times 0.001$ | $gxg_{distal}$ $\times 0.001$ |
|---|---|---|---|---|---|
| Alanine aminotransferase | 22 | 44388817 | **9.967** | -0.052 | **11.238** |
| Apolipoprotein B | 11 | 116648917 | **7.450** | 0.907 | **8.481** |
| C-reactive protein | 1 | 66257838 | **8.700** | 1.256 | 8.587 |
| Cholesterol | 11 | 116648917 | **9.002** | 0.996 | **9.453** |
| Hemoglobin A1c | 8 | 41542093 | **5.084** | 3.785 | **6.509** |
| Lipoprotein-A | 6 | 160560845 | **13.744** | **20.051** | **18.316** |
| | | 160578069 | **7.050** | **41.448** | **14.176** |
| | 19 | 45414399 | **9.567** | 0.781 | **11.421** |
| Mean platelet volume | 20 | 57597970 | **6.834** | 1.478 | **8.886** |
| Monocyte count | 13 | 28623048 | **3.855** | 2.204 | **7.772** |
| SHBG | 17 | 7145117 | **8.215** | **7.242** | **10.908** |
| | | 7254315 | **7.073** | **8.189** | **9.768** |
| Testosterone | 7 | 99032593 | **7.908** | 1.410 | **8.075** |
| | 12 | 2977954 | **8.692** | **3.437** | **9.192** |
| Triglycerides | 11 | 116648917 | **9.082** | 1.598 | **17.031** |
| Urate | 1 | 145630111 | **3.170** | **2.819** | **4.416** |
